# Likelihood-based Tests for Detecting Circadian Rhythmicity and Differential Circadian Patterns in Transcriptomic Applications

**DOI:** 10.1101/2021.02.23.432538

**Authors:** Haocheng Ding, Lingsong Meng, Andrew C. Liu, Michelle L. Gumz, Andrew J. Bryant, Colleen A. Mcclung, George C. Tseng, Karyn A. Esser, Zhiguang Huo

## Abstract

Circadian rhythmicity in transcriptomic profiles has been shown in many physiological processes, and the disruption of circadian patterns has been founded to associate with several diseases. In this paper, we developed a series of likelihood-based methods to detect (i) circadian rhythmicity (denoted as LR rhythmicity) and (ii) differential circadian patterns comparing two experimental conditions (denoted as LR diff). In terms of circadian rhythmicity detection, we demonstrated that our proposed LR rhythmicity could better control the type I error rate compared to existing methods under a wide variety of simulation settings. In terms of differential circadian patterns, we developed methods in detecting differential amplitude, differential phase, differential basal level, and differential fit, which also successfully controlled the type I error rate. In addition, we demonstrated that the proposed LR diff could achieve higher statistical power in detecting differential fit, compared to existing methods. The superior performance of LR rhythmicity and LR diff was demonstrated in two real data applications, including a brain aging data (gene expression microarray data of human postmortem brain) and a time-restricted feeding data (RNA sequencing data of human skeletal muscles). An R package for our methods is publicly available on GitHub https://github.com/diffCircadian/diffCircadian.

## Introduction

Circadian rhythms are an endogenous ∼ 24 hours cycle of behavior and physiology including sleep-wake cycles, body temperature, and melatonin [1, 2, 3, 4]. Underlying circadian rhythms is the clock mechanism that is found in virtually all cells of body. This mechanism is defined by a transcriptional-translational feedback loop involving a set of core clock genes [5, 6], including *CLOCK, BMAL1*, period family (*PER1, PER2, PER3*), and cryptochrome family (*CRY1, CRY2*). Beyond the core clock mechanism, genome-wide transcriptomic studies have uncovered circadian genes expression patterns in many tissues, including postmodern brain [7, 8], skeletal muscle [9], liver [10], and blood [11]. Zhang et al. [12] and Ruben et al. [13] conducted genome-wide circadian analyses using transcriptomic data of 12 unique mouse organs and 13 unique human organs, respectively, and showed that the profiles of circadian gene expression were tissue specific. It is now recognized from studies in humans and rodents that disruption in clock and circadian gene expression are linked to diseases including type II diabetes [14], sleep [11], major depression disease [15], aging [7], schizophrenia [8], and Alzheimer’s disease [16].

In the literature, several algorithms have been developed to detect circadian rhythmicity, including F-test via cosinor-based rhythmometry [17], Lomb-Scargle periodograms [18], COSOPT [19], ARSER [20], RAIN [21], JTK CYCLE [22], and MetaCycle [23]. These algorithms were widely applied in transcriptomic studies, and the comparisons of these algorithm have been evaluated in several review studies [24, 25, 26]. Though promising, concerns have been raised [25] that the p-values generated by many of these existing methods may not be correct (i.e., do not follow a uniform distribution [i.e., U(0, 1)] under the null), implying a potential inflated or deflated type I error rate.

Another increasingly important research question is to identify differential circadian patterns associated with different experimental conditions [27, 28, 11]. Figure 1 shows 4 types of differential circadian patterns identified in our brain aging data application (see Section 4.1 for details), among which 31 subjects were from the young group (age *≤* 40 years), and 37 subjects were from the old group (age *>* 60 years). Gene *CIART* in Figure 1a shows the differential amplitude, where the amplitude in the young group is larger than the old group; Gene *PER2* in Figure 1b shows the differential phase, where the phases in young and old groups are different; Gene *TRIB2* in Figure 1c shows the differential basal level, where the basal level in the young group is higher than the old group; Gene *MYO5A* in Figure 1d shows the differential fit, where there exists a good circadian rhythmicity fit in the young group, but not in the old group. The definition of amplitude, phase, and basal level is illustrated in Figure 2. The traditional approach to compare circadian rhythmicity between two experimental conditions is to adopt a hard threshold (e.g., *p ≤* 0.01) as the significance cutoff, and then declare deferential circadian rhythmicity if a gene is significant in only one condition [11, 29]. Though straightforward, this approach may fail under the following two scenarios. Scenario (i): gene *PER2* in Figure 1b is showing significant circadian rhythmicity in both the young group (*p* = 1.60 × 10^−4^) and the old group (*p* = 7.37× 10^−5^), and thus did not satisfy the definition of differential circadian pattern. However, Figure 1b shows a clear phase difference comparing young and old groups, and the underlying differential phase p-value using our proposed method was 5.43 × 10^−5^. Scenario (ii): gene *EEF2K* had a circadian p-value 0.0096 in young group, and a p-value 0.0305 in the old group. Though this gene satisfied this definition of differential circadian pattern using *p ≤* 0.01 as the significance criteria, the rhythmicity p-values under both conditions were close to 0.01. In fact, the resulting differential fit p-value using our proposed method was 0.709, indicating *EEF2K* was not showing differential circadian pattern comparing the young group and the old group.

**Fig. 1.**
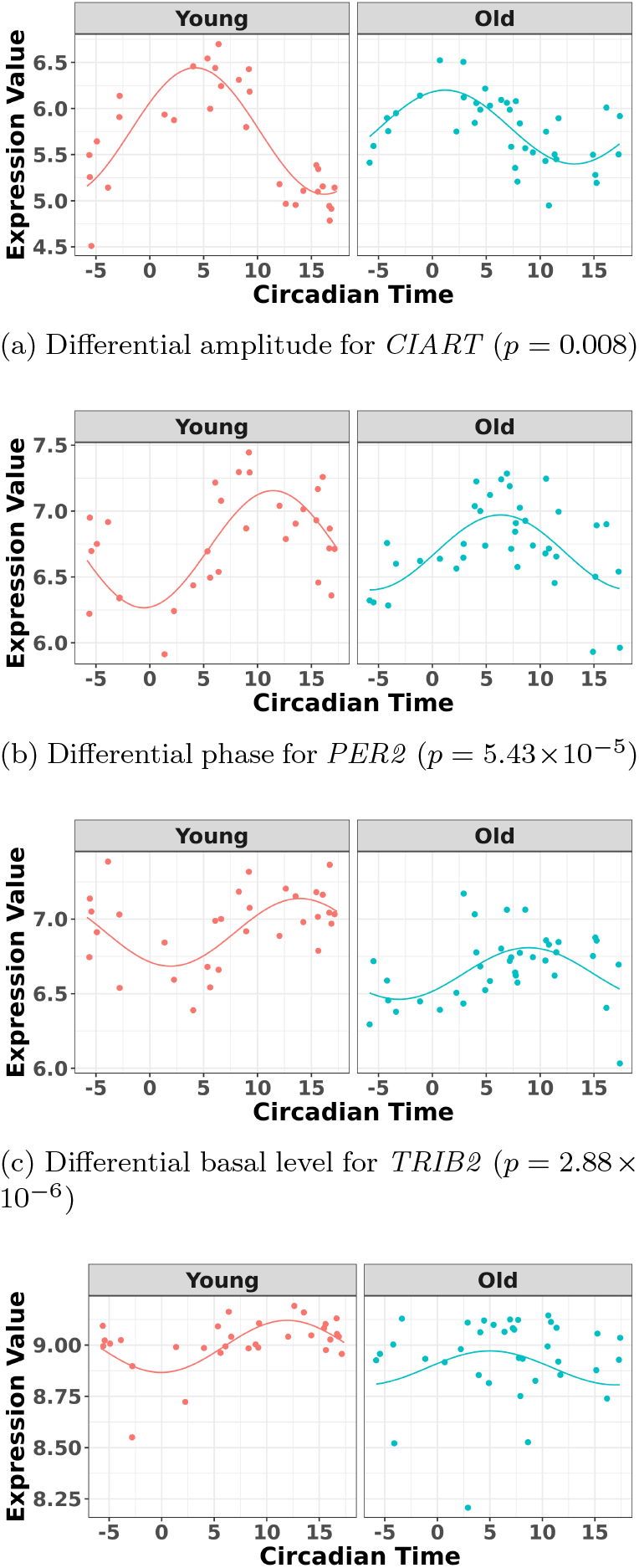
The most significant genes showing four types of differential circadian patterns from the brain aging data.

**Fig. 2.**
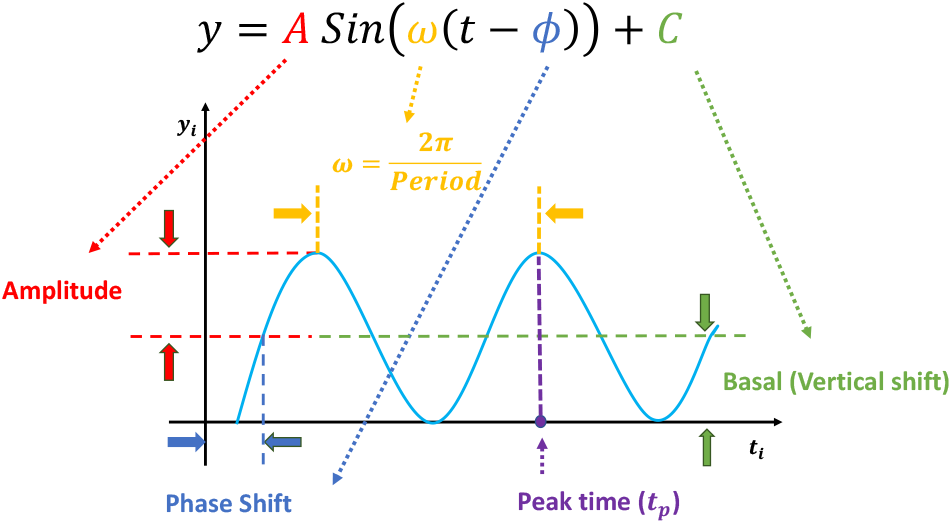
Illustration of a sinusoidal wave fitting and its related terminologies.

In the literature, some methods have been developed to identify genes showing differential circadian patterns. Chen et al. [7] developed a permutation test to quantify the statistical significance of these 4 types of differential circadian patterns. However, the non-parametric permutation test could suffer from low p-value precision and heavy computational burdens. DODR [30] and LimoRhyde [31] were developed to examine the hypothesis that the circadian rhythmicity across two conditions are identical, but they failed to further categorize different subclasses of differential circadian patterns illustrated in Figure 1. More recently, circaCompare [32] was developed to detect differential amplitude, differential phase, and differential basal level using non-linear least square methods, but it could not characterize differential fit. To our knowledge, there is still a lack of unified parametric method that could identify all 4 differential circadian patterns simultaneously. In addition, the performance of these existing methods has not been systematically evaluated.

In the statistics field, likelihood-based methods enjoyed tremendous popularity for its simplicity when testing single parameter and its flexibility to extend to test multiple parameters or complex models. In addition, the testing procedures based on the likelihood-based methods are generally considered as asymptotically the most efficient. However, this concept has not been fully developed in the field of circadian analysis. To close these research gaps, and to fully incorporate the merit of likelihood-based approaches, we propose a series of likelihood-based methods to detect circadian rhythmicity (within one condition) as well as differential circadian patterns (comparing two conditions). The contribution and novelty of this paper includes: (i) systematically evaluated the accuracy of p-values in detecting circadian rhythmicity of our likelihood-based methods and other existing methods. (ii) the first to propose likelihood-based methods to identify all 4 types of differential circadian patterns; (iii) systemically evaluated our likelihood-based methods in detecting differential circadian patterns, and compared with existing methods in terms of the correctness of p-value and statistical power; (vi) implemented our proposed methods in R software package, which has been made publicly available on GitHub.

## Method

We developed likelihood-based methods for (i) circadian rhythmicity detection within one experimental condition and (ii) differential circadian pattern analysis comparing two experimental conditions. The statistical inference of these methods were based on the Wald statistics and the likelihood ratio statistics. Since the accurate inference of the likelihood-based methods required large sample size, we also employed finite sample corrections to improve the performance under small sample sizes.

### Notations for a sinusoidal wave fitting

Our methods assume that the relationship between the gene expression level and the circadian time fits a sinusoidal wave curve. As illustrated in Figure 2, denote *y* as the expression value for a gene; *t* as the circadian time; *C* as the basal level (vertical shift of the sinusoidal wave baseline from 0); *A* as the amplitude. *ω* is the frequency of the sinusoidal wave, where 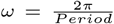. Without loss of generality, we set *period* = 24 hours to mimic the diurnal period. *ϕ* is the phase of the sinusoidal wave curve. Whenever there is no ambiguity, we will omit the unit “hours” in period, phase, and other related quantities. Due to the periodicity of a sinusoidal wave, (*ϕ*_1_, *ϕ*_2_) are not identifiable when *ϕ*_1_ = *ϕ*_2_ + 24. Therefore, we will restrict *ϕ ∈* [−6, 18). *ϕ* may be difficult to read from a sinusoidal wave (See Figure 2), and a closely related quantify is the peak time *t*_*P*_. The connection between *ϕ* and *t*_*P*_ is that *ϕ* +*t*_*P*_ = 6 *±*24*N*, where *N* is an arbitrary natural number.

### Circadian Rhythmicity Detection

In this section, we develop likelihood-based methods to test the existence of a circadian rhythmicity within one experimental condition. Denote *y*_*i*_ is the expression value of one gene for subject *i*(1 *≤ i ≤ n*), where *n* is the total number of subjects. *t*_*i*_ is the circadian time for subject *i*. We assume:

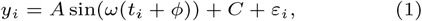

where *ε*_*i*_ is the error term for subject *i*; we assume *ε*_*i*_’s are identically and independently distributed (i.e., *iid*) and *ε*_*i*_ ∼ *N*(0, *σ*^2^), where *σ* is the noise level. To benchmark the goodness of sinusoidal wave fitting, we define the coefficient of determination 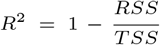, where 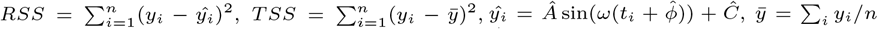, with *Â*, 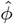, and *Ĉ* being the fitted value for *A, ϕ*, and *C* in Equation 1 under least square loss, respectively. *R*^2^ ranges from 0 to 1, with 1 indicating perfect sinusoidal wave fitting, and 0 indicating no fitting at all. Based on these assumptions, we derive procedures for testing circadian rhythmicity. For the ease of discussion, we re-write Equation 1 as

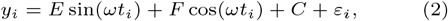

where *E* = *A* cos(*ωϕ*), and *F* = *A* sin(*ωϕ*). The hypothesis setting for testing the existence circadian rhythmicity is *H*_0_: *E* = *F* = 0 *v*.*s. H*_*A*_: *E* ≠ 0 or *F* ≠ 0. We will derive the Wald statistics and the likelihood ratio statistics to perform hypothesis testing. Since both Wald statistics and likelihood ratio statistics are designed based on large sample theories, we will also employ finite sample statistics for these methods.

### Likelihood Ratio Test

Based on Equation 2, the likelihood function of all *n* samples is:

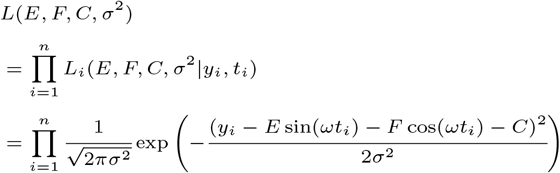

The log-likelihood function is:

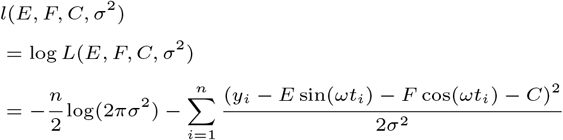

Under 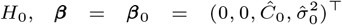, and 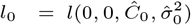, where ***β***_0_ is the least square estimate of Equation 2 under *H*_0_. Under 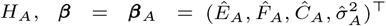 and 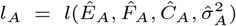, where ***β***_*A*_ is the least square estimate of Equation 2. The likelihood ratio test statistic is: *T*^*LR*^ = −2(*l*_0_ − *l*_*A*_). Since the degree of freedom is 2, under *H*_0_, 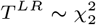.

### Wald Test

The Wald test statistic can be derived as *T*^*W ald*^ = (***β***_*A*_ − ***β***_0_)^T^ *I*(***β***_*A*_)(***β***_*A*_ − ***β***_0_), where *I*(***β***_*A*_) is Fisher information matrix evaluated at ***β***_*A*_. Under *H*_0_, 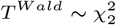.

### Finite Sample Wald/LR Tests

The Wald test and the likelihood ratio test may have inflated type I error when sample size is small since they rely on large sample asymptotic theory. Parker [33] introduced finite sample Wald and likelihood ratio test statistics, which could better control the type I error rate to the nominal level even with small sample sizes. The finite sample Wald statistics 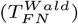 and the finite sample likelihood ratio statistics 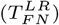 can be derived as the following:

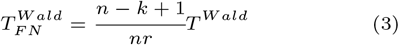

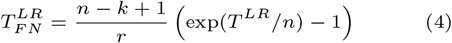

Where *k* = 4 is total number of parameters and *r* = 2 is number of parameters of interest. Under the null hypothesis, 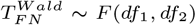, and 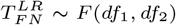, where *df*_1_ = *r* and *df*_2_ = *n* − *k* + 1.

### F Test

The F-test method to detect the circadian rythmicity has been previously established [17]. F test constructs its test statistic by decomposing total variability into model sum of square, and residual sum of square, which is closely related to our proposed finite sample likelihood method. Thus, we also describe the F-test method in our manuscript, and will draw connection between F-test and our likelihood method.

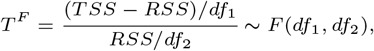

where residual sum of squares 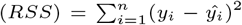, total sum of squares 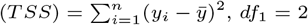 and *df*_2_ = *n* − 3. Under the null hypothesis, *T*^*F*^ ∼ *F* (*df*_1_, *df*_2_).

### Other competing methods

We will compare our proposed likelihood-based method to other existing methods, including F-test [17], ARSER [20], Lomb-Scargle periodograms [18], JTK CYCLE [22], RAIN [21], and the permutation test [7]. ARSER, RAIN, and JTK CYCLE have some special requirement for the input circadian time – the input circadian time has to be integer value, and the intervals between two adjacent circadian time points must be the same. Thus, we will accommodate such design in our simulation settings when needed.

### Differential Circadian Analysis

In this section, we develop likelihood-based testing procedures to identify genes showing differential circadian patterns, including (a) differential amplitude, (b) differential phase, (c) differential basal level, and (d) differential fit, as shown in Figure 1.

Denote *y*_1*i*_ as the gene expression value of subject *i*(1 *≤ i ≤ n*_1_) in experimental condition 1, where *n*_1_ is the total number of subjects; *t*_1*i*_ is the circadian time for subject *i. y*_2*j*_ is the gene expression value of subject *j*(1 *≤ j ≤ n*_2_) in experimental condition 2, where *n*_2_ is the total number of subjects; *t*_2*j*_ is the circadian time for subject *j*. Note that *y*_1*i*_ and *y*_2*j*_ are from the same gene, but under different experimental conditions. We assume the following models:

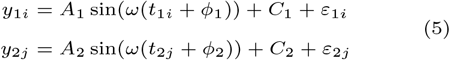

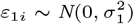 is the error term for subject *i* (1 *≤ i ≤ n*_1_) for experimental condition 1 and 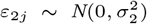 is the error term for subject *j* (1 *≤ j ≤ n*_2_) for experimental condition 2. These error terms are assumed to be *iid. A*_1_, *ϕ*_1_, *C*_1_, and 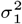 are the amplitude, phase, basal level, and noise level for the experimental condition 1, and *A*_2_, *ϕ*_2_, *C*_2_, and 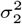 are for experimental condition 2.

### Hypothesis testing framework for differential circadian analysis

Below we state the null hypothesis and the alternative hypothesis for testing these four categories of differential circadian patterns, based on Equation 5.

1. Differential amplitude: *H*_0_: *A*_1_ = *A*_2_ = *A*_*c*_ *v*.*s. H*_*A*_: *A*_1_ ≠*A*_2_.
2. Differential phase: *H*_0_: *ϕ*_1_ = *ϕ*_2_ = *ϕ*_*c*_ *v*.*s. H*_*A*_: *ϕ*_1_ ≠ *ϕ*_2_.
3. Differential basal level: *H*_0_: *C*_1_ = *C*_2_ = *C*_*c*_ *v*.*s. H*_*A*_: *C*_1_ ≠*C*_2_.
4. Differential fit: 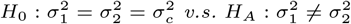.

#### Remark 1

*(i) As suggested by Chen et al. [7], the circadian rhythmicity can be characterized by the goodness of fit statistics R*^2^. *Since it is not easy to derive statistical inference on R*^2^, *we will use a closely related quantity, σ*^2^, *to quantify the goodness of fit. (ii) The prerequisite for differential amplitude, differential phase, and differential basal level is that there should exist circadian rhythmicity in both conditions under comparisons. Therefore, we suggested users to set p ≤* 0.01 *or p ≤* 0.05 *from our previous likelihood-based circadian rhythmicity test to ensure the existence of the circadian rhythmicity in both conditions. (iii) The prerequisite for differential fit is that there should exist a circadian rhythmicity in either experimental conditions. We suggested users to set p ≤* 0.01 *or p ≤* 0.05 *from our previous likelihood-based circadian rhythmicity test to ensure such prerequisite*.

### Likelihood ratio test

Based on Equation 5, the log-likelihood function for *n*_1_+*n*_2_ samples in both experimental conditions is:

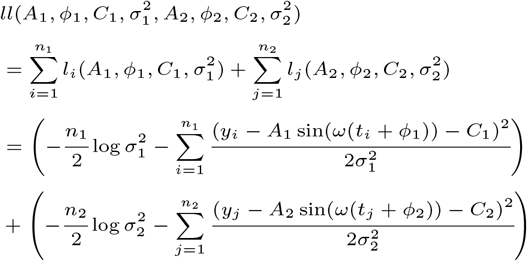

The test statistic is: *D*^*LR*^ = −2(*ll*_0_ − *ll*_*A*_), where *ll*_0_ is the log likelihood under *H*_0_; and *ll*_*A*_ is the log likelihood under *H*_*A*_. Here the null can be one of the following: (a) 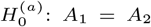 for differential amplitude; (b) 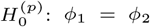 for differential phase; (c) 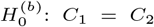 for differential basal level; (d) 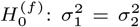 for differential fit. For all these null hypotheses, the degree of freedom is 1, and 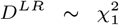 under *H*_0_. For example when testing different amplitude, under 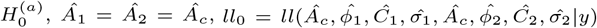; and under 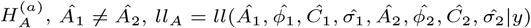.

### Wald Test

Denote 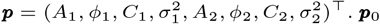. ***p***_0_ is ***p*** under *H*_0_, where *H*_0_ is one of the null hypotheses in Section 2.3.1; ***p***_1_ is ***p*** under *H*_*A*_, where there is no restriction on ***p***. Then the Wald test statistic is *D*^*W ald*^ = (***p***_1_ − ***p***_0_)^T^ *I*(***p***_1_)(***p***_1_ −***p***_0_). Under *H*_0_, 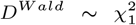, where *I*(***p***_1_) is Fisher information matrix evaluated at ***p***_1_.

### Finite Sample Wald/LR Tests

Again, in order to control type I error for small sample sizes, we derive finite sample version of the Wald statsitics and likelihood ratio statistics 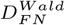 and 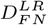 by Equations 3 and 4. Under *H*_0_, 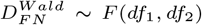, and 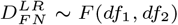, where *df*_1_ = *r* and *df*_2_ = *n*−*k* +1.

### Competing methods for differential circadian analysis

We will compare the performance of our method with other existing methods, including the permutation test [7], DODR [30], LimoRhyde [31], and circaCompare [32]. We acknowledge that HANOVA, robustDODR and LimoRhyde are designed to detect differential rhythmicity (i.e., whether the circadian rhythmicity across two conditions are identical), and cannot distinguish the 4 subcategories in Figure 1. Thus we will apply these two methods in detecting differential fit, which is closely related to differential rhythmicity conceptually; circaCompare can examine differential amplitude, differential phase, and differential basal level; while the permutation test as well as our proposed method can examine all four types if differential circadian patterns illustrated in Figure 1.

## Simulation

In terms of circadian rhythmicity detection, we demonstrated that our proposed method correctly controlled the type I error rate to the nominal level, while some of the other methods failed to control the type I error rate. In terms of differential circadian pattern detection, our method still controlled the type I error rate to the nominal level. For differential fit, which is one type of the differential circadian pattern shown in Figure 1d, we demonstrated our method achieved higher statistical power compared to the existing methods.

### Simulation for Circadian Rhythmicity Analysis

#### Simulation Settings

Denote *i*(1 *≤ i ≤ n*) as the sample index, where *n* was the total number of samples. The circadian time *t*_*i*_ for sample *i* was generated from uniform distribution UNIF(0, 24). We simulated the gene expression value for sample *i* using Equation 1.

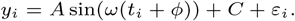

Our basic parameter setting for simulation is listed as below. For each gene, the sample size *n* was set to be 12; the circadian time were sampled every 2 hours (i.e., *t*_1_ = 1, *t*_2_ = 3,…, *t*_12_ = 23), such integer circadian time and evenly spaced interval time are required by some other existing methods. Whenever the statistical methods have no such requirement, we sampled circadian time directly from UNIF(0, 24). Amplitude *A* was fixed at 1; phase *ϕ* was generated from UNIF(0, 24). Basal level C was generated from UNIF(0, 3). Error term *ε*_*i*_ was generated from normal distribution *N* (0, *σ*^2^) where *σ*^2^ was set to be 1. We simulated *G* = 10, 000 genes for each simulation, and each simulation was repeated *B* = 10 times to increase numbers of replications and to obtain an standard error estimate. To examine whether our method is robust against higher signal-noise ratios, correlated gene structures, and violations of normality distributions, we further simulates the following variations:

1. Impact of sample sizes. We varied *n* = 6, 12, 24, 48, 96 while fixing other parameters in the basic parameter setting fixed. Note that when *n >* 24, we would allow repetitive circadian time for different samples. For example, when *n* = 48, the circadian time sequence would be *t*_1_ = 1, *t*_2_ = 1, *t*_3_ = 2,…, *t*_47_ = 24, and *t*_48_ = 24.
2. Impact of signal noise ratio. The signal noise ratio is defined as 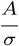. Thus we varied *σ* = 1, 2, 3 to mimic varying levels of signal noise ratio, while fixing other parameters in the basic parameter setting.
3. Impact of correlated genes. In transcriptomic data applications, individual genes are can be be correlated. Thus, we simulated the following correlated structure. For every m = 50 genes, we simulated

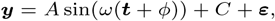

where ***y*** = (*y*_1_,…, *y*_*m*_), ***t*** = (*t*_1_,…, *t*_*m*_), and ***ε*** = (*ε*_1_,…, *ε*_*m*_). In this case, ***ε*** were generated from a multivariate normal distribution *MV N* (0, **σ**). And **σ** *∈* ℝ^*m* × *m*^ was the covariance matrix generated from the inverse Wishart distribution *W* ^−1^(**ϕ**, *v*). In order to mimic correlated gene structure, we first designed ϕ ′ = (1−*ρ*)*I*_*m*× *m*_ +*ρJ*_*m* × *m*_, and then standardized ϕ to correlation matrix ϕ, where *I*_*m* × *m*_ was the identify matrix, and *J*_*m* × *m*_ *∈* ℝ^*m* × *m*^ a matrix with all elements 1. We fixed *v* to be 60, and vary *ρ* = 0, 0.25, 0.5.
4. Violation of the Gaussian assumption. Instead of assuming the error term was generated from a standard normal distribution (i.e., *N* (0, 1)), we generated *ε*_1_ ∼ *t*(3), *t*(5), *t*(10), *t*(∞), where *t*(*df*) is the t-distribution with degree of freedom *df*. This family of t-distributions represents long tailed error distribution, with smaller *df* indicating longer tailed error distribution, and thus larger violation of the normality assumption. When *df* = ∞, *t*(∞) is the same as *N* (0, 1).

#### The best performer of the likelihood based methods in detecting circadian rhythmicity

Before comparing with other existing methods, we first evaluated the Type I error rate (nominal *α* level 5%) of our proposed four likelihood-based methods in detecting circadian rhythmicity, including Wald test (regular), Wald test (finite sample), likelihood ratio test (regular), likelihood ratio test (finite sample). Since the limiting distribution of both Wald statistics (finite sample) and likelihood ratio statistics (finite sample) follows an F distribution, we also include the F-test method [17] as benchmark.

Figure S1 showed type I error rates (nominal *α* level 5%) of our proposed four methods and the F-test method. Regardless of the varying sample sizes, the Wald test (finite sample), the likelihood ratio test (finite sample), and the F-test controlled the type I error rate close to the 5% nominal level, while the Wald test (regular) and the likelihood ratio test (regular) obtained inflated type I error rate. The Wald test (regular) and the likelihood ratio test (regular) had better performance when sample size became larger, which was not unexpected because these asymptotic tests rely on large sample sizes. Remarkably, we observed that the Wald test (finite sample) and the likelihood ratio test (finite sample) achieved almost the same test statistics as the F-test, indicating the finite sample approximation procedure [33] successfully convert our likelihood-based statistics to F statistics.

Similar results was also observed by varying signal noise ratio (Figure S2) and varying the strength of gene correlations (Figure S3). The Wald test (finite sample), the likelihood ratio test (finite sample), and the F-test could better control the type I error rate to the 5% nominal level compared to the the Wald test (regular) and the likelihood ratio test (regular).

As shown in Figure S4, when we varied the level of normality violation by varying *df* of the t-distribution, we observed that all test procedures became more conservative, which is not unexpected because our likelihood-based method assumed Gaussian error in the likelihood construction. In practice, if the residuals (i.e., *y*_*i*_ − *ŷ*_*i*_) violated the Gaussian distribution, we would recommend data transformations (e.g., log transformation) to improve normality.

To summarize, the Wald test (finite sample) and the likelihood ratio test (finite sample) are the best performer of our proposed likelihood-based methods in detecting circadian rhythmicity, which could control the type I error rate to the nominal level under the Gaussian assumption. And these two methods are equivalent to the F-test method in terms of the test statistics. Therefore, we will pick up the likelihood ratio test (finite sample) as the representative of our proposed methods in detecting circadian rhythmicity, and we denoted LR rhythmicity as the short name for this method in all later evaluations.

#### Type I Error rate comparison with other methods

We compared the likelihood-based method (LR rhythmicity) with other existing methods in detecting circadian rhythmicity, including Lomb-Scargle, JTK, ARSER, Rain, and permutation. We excluded the F test in our evaluation, since it is essentially the same as LR rhythmicity. Figure 3 showed type I error rates by varying sample sizes. In general, LR rhythmicity and the permutation test controlled the type I error rate to the 5% nominal level, while the other methods had inflated or conservative type I error rate. Similar results was also observed by varying signal noise ratio (Figure S5) and varying the strength of gene correlations (Figure S6). As shown in Figure S7, we observed that a larger violation of normality assumption will lead to more conservative type I error rate for LR rhythmicity, while the permutation test can still control the type I error rate to the 5% nominal level.

**Fig. 3.**
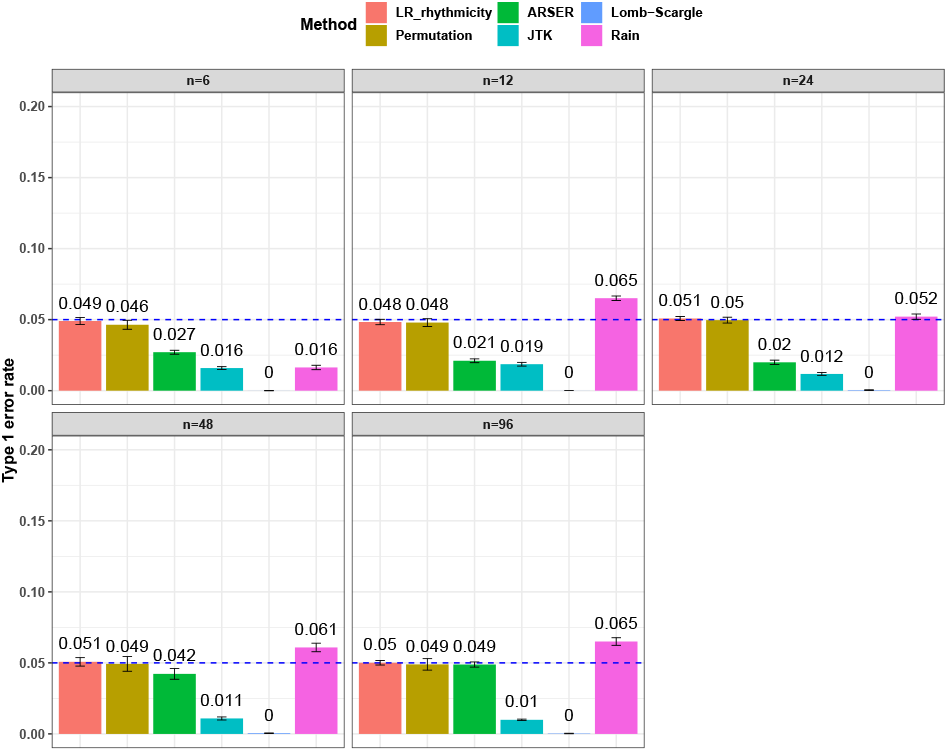
Type I error rate at nominal *α* level 5% for 6 different methods in detecting circadian rhythmicity. The sample sizes were varied at n=6, 12, 24, 48, and 96. The blue dashed line is the 5% nominal level. A higher than 5% blue dashed line bar indicates an inflated type I error rate; a lower than 5% blue dashed line bar indicates a conservative type I error rate; and a bar at the blue dashed line indicates an accurate type I error rate (i.e., *p* = 0.05). The standard deviation of the mean type I error rate was also marked on the bar plot.

To summarize, under the Gaussian assumption (i.e., the residual follows normal distribution), only the LR rhythmicity and the permutation test can achieve nominal type I error rate control (i.e., 5%).

### Power analysis

For the power analysis, we only examined the method that could successfully control the type I error rate to the 5% nominal level. Otherwise the power is directly not comparable because it cannot be distinguished whether a higher/lower power is a result of the test procedure itself, or because of inflated/conservative type I error rate control. Only the LR rhythmicity and the permutation test survived these criteria. Figure S8 shows the power with respect to varying sample sizes. Both these methods are similarly powerful at 5% nominal level of type I error rate. When the sample size is larger, both tests became more powerful. However, we want to point out that the precision of the permutation test depends heavily on the number of permutations. For example, it may need at least 1,000,000 permutations in order to achieve a *p <* 10^−6^, which could be a computational burden. The LR rhythmicity has no such restriction, and could obtain an arbitrarily small p-values without extra computational concerns.

### Differential Circadian Analysis

In this section, we used simulation to evaluate the performance of the likelihood-based method in detecting differential circadian patterns, including differential amplitude, differential phase, differential basal level, and differential fit. We first compared among our proposed likelihood-based methods including Wald test (regular), Wald test (finite sample), likelihood ratio test (regular), likelihood ratio test (finite sample). We found that likelihood ratio test (finite sample) was the best performer of our proposed methods. We then compared this best performer with other existing methods for differential circadian pattern analysis, including Circacompare, limorhyde, HANOVA, robustDODR, and the permutation test under variety of simulation settings.

### Simulation settings

The simulation setting is based on Equation 5. The basic parameter setting for simulation is listed as below. We set number of genes *G* = 10, 000 and the sample size *n* was set to be 10. For each gene *g* (1 *≤ g ≤ G*), amplitudes *A*_1_ = *A*_2_ were set to be 3; phases *ϕ*_1_ = *ϕ*_2_ were generated from UNIF(0, 24). Basal levels *C*_1_ = *C*_2_ were generated from UNIF(10, 13). Error terms *ε*_*i*_, *ε*_*j*_ were generated from normal distribution 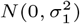 and 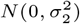, respectively. 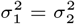 were set to be 1. This simulation was repeated 10 times to increase numbers of replications and to obtain standard error estimate. To examine the impact of sample size, correlation between genes, and distribution violations, we further simulated the following variations.

1. Impact of sample sizes. We varied *n* = 10, 20, 50 while fixing other parameters in the basic parameter setting.
2. Impact of correlated genes. For every m = 50 genes, we simulated the correlated gene structure as described in Section 3.1.1. We varied the strength of correlation *ρ* = 0, 0.25, 0.5 while fixing other parameters in the basic parameter setting.
3. Violation of the Gaussian assumption. As described in Section 3.1.1, we varied the error distribution *ε*_1_ = *ε*_2_ ∼ *t*(3), *t*(5), *t*(10), *t*(∞) to mimic different levels of violation of normality assumptions.

### The best performer of the likelihood-based methods in detecting differential circadian patterns

We evaluated the type I error rate (nominal *α* level 5%) of our proposed likelihood-based methods, including Wald test (regular), Wald test (finite sample), likelihood ratio test (regular), likelihood ratio test (finite sample), under all pre-mentioned simulation settings. Figure S9, S10 show that the likelihood ratio test (finite sample) had the best performance in terms of type I error rate control with varying sample size or strength of correlation among genes. Thus, we denoted this method as LR diff, and will further compared LR diff with other existing methods. In Figure S11, when there was a violation of the Gaussian assumption, we observed that LR diff still controlled the type I error rate for differential amplitude, differential phase, and differential basal levels, but resulted in inflated type I error rate of differential fit. This is not unexpected since likelihood-based methods utilized the Gaussian assumption to derive the test statistics. Under this situation, we would recommend users to take transformation (i.e., log transformation) to improve normality (See Section 5 for more discussions).

### Type I Error rate comparison with other methods

We evaluated the type I error rate (nominal *α* level 5%) of the following methods: LR diff, Circacompare, limorhyde, HANOVA, robustDODR, and permutation test under different simulation settings (See Section 3.2.1 for details). Here, LR diff and the permutation test are applicable for testing all 4 types of differential circadian analysis in Figure 1; HANOVA, robustDODR and LimoRhyde are designed to detect differential rhythmicity (i.e., whether the circadian rhythmicity across two conditions are identical), and cannot distinguish the 4 subcategories.

Thus we will apply these three methods in detecting differential fit, which is closely related to differential rhythmicity conceptually; and Circacompare is applicable for testing differential amplitude, differential phase, and differential basal levels (Also see Table 1 for for their applicability).

**Table 1.** Comparison of LR diff with other existing methods in detecting differential circadian patterns. ✓ indicates a method is applicable; or could control the type I error to the nominal level; * indicates the most powerful method among all applicable methods. − indicates the method could roughly control the type I error to the nominal level, but with a non-negligible deviation.

|  | Differential amp/phase/basal |  | Differential fit |  |
| --- | --- | --- | --- | --- |
|  | Applicable | Type I error | Applicable | Type I error |
| LR_diff | ✓ | ✓* | ✓ | ✓* |
| Permutation | ✓ | ✓* | ✓ | ✓ |
| Circacompare | ✓ | ✓* |  |  |
| limorhyde |  |  | ✓ | ✓ |
| HANOVA |  |  | ✓ | – |
| robustDODR |  |  | ✓ | – |

1. Impact of sample sizes. Based on the basic parameter setting, we varied *n* = 10, 20, 50. Figure 4 shows the type I error rate control for the 6 methods. Among which 3 methods were applicable for detecting differential amplitude, differential basal level, differential phase, including LR diff, Circacompare and the permutation test. All these methods could control the type I error rate to the 5% nominal level, though Circacompare is slightly better than LR diff and the permutation test. In addition, 5 methods were applicable for detecting differential fit, including LR diff, limorhyde, HANOVA, robustDODR, and the permutation test. We observed that LR diff, limorhyde, and the permutation test could control the type I error rate to the 5% nominal level, while HANOVA and robustDODR may have slightly inflated type I error rate. We observed their performance did not rely heavily on the sample size, which is expected since these methods did not necessarily rely on large sample size.
2. Impact of correlated genes. Figure S12 shows the type I error rate control by varying the strength of correlations between genes. Similar to the previous simulation setting, we did not observe the correlated gene structure had a big impact on their performance.
3. Violation of the Gaussian assumption. Instead of assuming the error term was generated from a standard normal distribution (i.e., *N* (0, 1)), we generated *ε*_1_ ∼ *t*(3), *t*(5), *t*(10), *t*(*∞*), where *t*(*df*) is the t-distribution with degree of freedom *df*. Smaller *df* represents longer tailed error distribution, and thus larger violation of the normality assumption. Figure S13 shows the type I error rate control for the 6 methods. In terms of differential amplitude, differential basal level, differential phase, LR diff, Circacompare and the permutation test successfully controlled the type I error rate to the 5% nominal. In terms of differential fit, we observed that the LR diff would obtain inflated type I error rate, while the performance of limorhyde, HANOVA, robustDODR, and the permutation test were similar regardless of violation of the Gaussian assumption. This is not unexpected because our likelihood-based method relied on the Gaussian assumption to derive its test statistics. Under this situation, we would recommend uses to take transformation (i.e., log transformation) to improve normality (See Section 5 for more discussions).

**Fig. 4.**
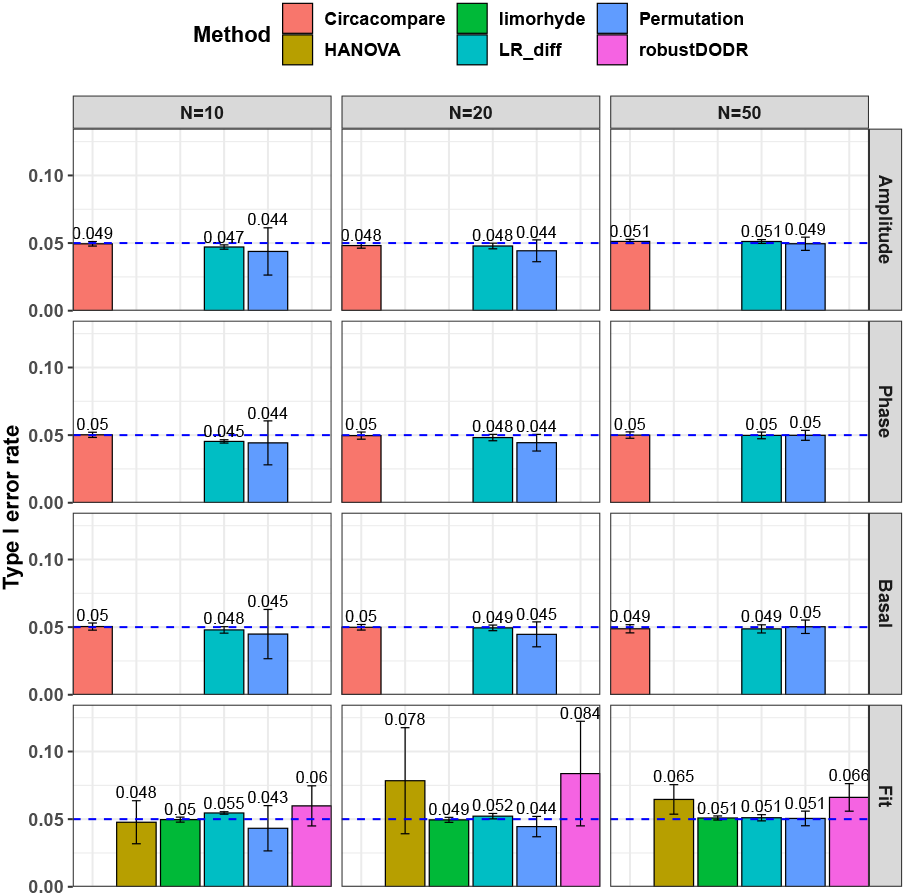
Type I error rate at nominal *α* level 5% for 6 different methods in detecting differential circadian patterns. The differential circadian patterns include differential amplitude (Amplitude), differential phase (Phase), differential basal level (Basal), and differential fit (Fit). The sample sizes were varied at N=10, 20, and 50. The blue dashed line is the 5% nominal level. A higher than 5% blue dashed line bar indicates an inflated type I error rate; a lower than 5% blue dashed line bar indicates a conservative type I error rate; and a bar at the blue dashed line indicates an accurate type I error rate (i.e., p-value = 0.05). The standard deviation of the mean type I error rate was also marked on the bar plot.

To summarize, in terms of differential amplitude, differential basal level, differential phase, LR diff, Circacompare and the permutation test could control the type I error rate to the 5% nominal level. In terms of differential fit and under normality assumption, LR diff, limorhyde, and the permutation test could control the type I error rate to the 5% nominal level, while HANOVA and robustDODR may have slightly inflated type I error rate.

### Power analysis

In principle, all methods could control the type I error rate to the 5% nominal level, we included all these methods in the power evaluation (Figure 5). In terms of differential amplitude, differential basal level, differential phase, with increasing sample size or larger effect size, all three methods, including LR diff, Circacompare, and the permutation test became more powerful. Fixing the sample size and effect size, we observed that LR diff and Circacompare are a little bit more powerful than the permutation test. In terms of differential fit, remarkable, our proposed LR diff is much more powerful than the permutation test, limorhyde, HANOVA, and robustDODR. In addition, with increasing sample size or larger effect size, LR diff and the permutation test are becoming more powerful, while the other methods remained similar power or had a little bit elevated power. Table 1 summarizes and applicability, performance of type I error rate control and power for all these methods.

**Fig. 5.**
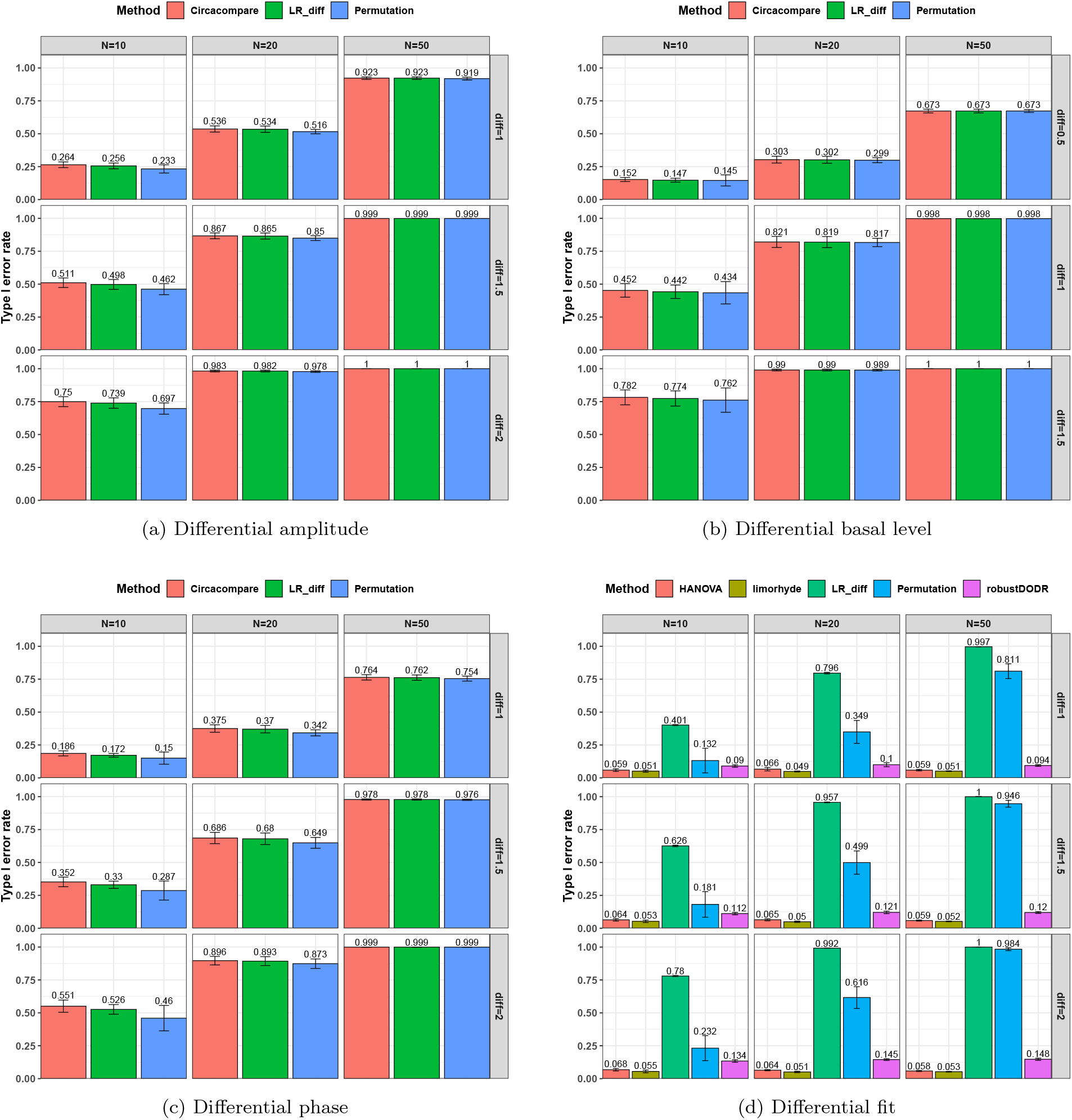
Power evaluation for 6 different methods in detecting differential circadian patterns. The differential circadian patterns include differential amplitude, differential phase, differential basal level, and differential fit. The sample sizes were varied at N=10, 20, and 50. The standard deviation of the mean type I error rate was also marked on the bar plot.

#### Remark 2

*We observed that our proposed LR diff had very similar type I error rate and statistical power compared to Circacompare. In fact, both LR diff and Circacompare were designed to address the same question (i*.*e*., *differential amplitude, phase, and basal level) by deploying cosinor-based rhythmometry. The difference is that LR diff utilized a likelihood ratio test, whereas Circacompare employed a non-linear least square approach. In addition, LR diff is capable of testing different fit, while Circacompare cannot be used to perform this test*.

## Real data applications

We evaluated our likelihood-based methods (LR rhythmicity and LR diff) in two real data applications, including a gene expression microarray data of human postmortem brain (comparing chronological age [i.e., young *v*.*s*. old]), and a gene expression RNA sequencing data of human skeletal muscles (comparing time-restricted feeding [i.e., restricted *v*.*s*. unrestricted]). Throughout this section, we used *p ≤* 0.01 as the cutoff to declare statistical significance unless otherwise specified. Since our likelihood-based method includes both circadian rhythmicity p-values and differential circadian pattern p-values, we denote *p*_*c*_ as a p-value for circadian rhythmicity detection (i.e., from LR rhythmicity), and *p*_*d*_ as a p-value for differential circadian pattern analysis i.e., from LR diff). *p*_*d*_ can also be expanded as 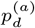 (differential amplitude); 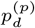 (differential phase); 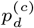 (differential basal level); 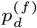 (differential fit). We did not compare our methods with existing methods in the real data application, because there is no underlying truth in the real data, and thus it is difficult to benchmark their performance.

### Brain aging data

We first examined our methods in a transcriptomic profile in a human postmortem brain data (Brodmann’s area 11 in the prefrontal cortex). Detailed description of this study has been previously described by Chen et al. [7]. The final samples included 146 individuals whose time of death (TOD) could be precisely determined. The mean age at death was 50.7 years; 78% of the individuals were male, and the mean postmortem interval was 17.3 hours. The TODs were further adjusted as the Zeitgeber time (ZT), which adjusted factors including time zone, latitude, longitude, and altitude. The ZT was used as the circadian time, which was comparable across all individuals. 33,297 gene probes were available in this microarray data, which was publicly available in GEO (GSE71620). After filtering 50% gene probes with lower mean expression level, 16,648 gene probes were kept.

### Circadian rhythmicity detection

Under *p*_*c*_ *≤* 0.01, we detected 528 significant circadian genes using LR rhythmicity. Figure 6 shows the 6 core circadian genes, including *PER1, PER2, PER3, ARNTL, NR1D1*, and *DBP*, which are known to have persistent circadian rhythmicity. All these 6 circadian genes rendered significant p-value (5.15 × 10^−24^ ∼ 3.59 × 10^−6^), showing the good detection power of our method in identifying circadian patterns. We further performed pathway enrichment analysis. Using pathway analysis *p ≤* 0.01 as cutoff, LR rhythmicity detected 4 pathways. The most significant pathway was the circadian rhythm signaling pathway (*p* = 3.16 × 10^−6^). The second most significant pathway was the senescence pathway (*p* = 4.27 × 10^−4^), which was also known to be associated with circadian oscillation [34].

**Fig. 6.**
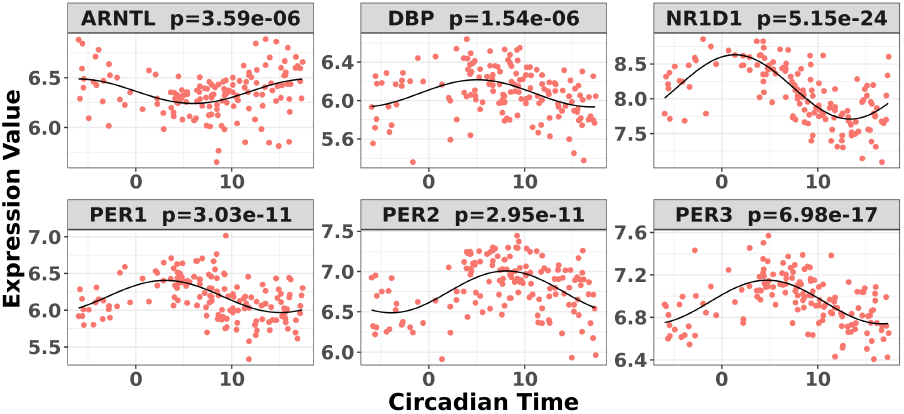
Circadian rhythmicity for 6 core circadian genes in the brain aging data, including *PER1, PER2, PER3, ARNTL, NR1D1*, and *DBP*, using LR rhythmicity.

### Differential circadian analysis

In order to examine whether the chronological age was associated with disruption of circadian patterns, we further performed differential circadian analysis comparing the young group and the old group using our likelihood-based method. We first divided the 146 individuals into two groups: young group (age *≤* 40, n=31) and old group (age *>* 60, n=37). Under *p*_*c*_ *≤* 0.01, we identified 205 genes showing circadian rhythmicity in young group and 164 genes in old group, with a total of 363 unique genes, and 6 common genes.

In terms of differential fit, we started with 363 candidate genes that showed circadian rhythmicity (*p*_*c*_ *≤* 0.01) in either young or old. Comparing the old group to the young group (baseline group), LR diff identified 6 genes showing differential fit 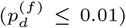. As shown in Figure 1d, *MYO5A* is the gene showing most differential fit 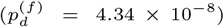, where there was circadian rhythmicity in the young group, but not in the old group. In terms of differential amplitude, differential phase, and differential basal level, we started with 6 candidate genes that showed circadian rhythmicity (*p*_*c*_ *≤* 0.01) in both young and old groups. Comparing the old group to the young group (baseline group), our likelihood-based method identified 1 gene showing differential amplitude 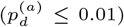, 4 genes showing differential phase 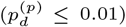, 2 genes showing differential basal level 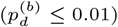. Figure 1a-1c showed the most significant genes in terms of differential amplitude (*CIART, p* = 0.008), differential phase (*PER2, p* = 5.43 × 10^−5^), and basal level (*TRIB2, p* = 2.88 × 10^−6^) comparing young and old groups, respectively.

Due to the small sample size and relatively weak transcriptomic alterations in brain tissues, the number of candidate genes for differential circadian analysis was small. Thus we further relaxed the criteria to be *p*_*c*_ *≤* 0.05, and we identified 897 rhythmic genes in the young group and 846 rhythmic genes in the old group. In terms of differential fit, among 1,688 genes that showed circadian rhythmicity (*p*_*c*_ *≤* 0.05) in either young or old group, LR diff identified 345 genes showing gain or loss of rhythmicity. In terms of differential amplitude, differential phase, and differential basal level, we started with 55 candidate genes that showed circadian rhythmicity (*p*_*c*_ *≤* 0.05) in both young and old groups. Comparing the old group to the young group (baseline group), LR diff identified 2 genes showing differential amplitude, 23 genes showing differential phase, 19 genes showing differential basal level.

### Time-restricted feeding data

We evaluated the performance of our likelihood-based methods in transcriptomic profiles of skeletal muscle tissue. 11 overweight or obese men were included in this dataset; the age range was 30-45 years; the body mass index [BMI] range was 27–35 *kg/m*^2^. These participants were randomized into time-restricted feeding (TRF) group and the un-restricted feeding (URF) group by adopting a cross over design, where each participant was assigned to both TRF and URF groups in different time periods. The skeletal muscle samples of each participant under each experimental group were repeatedly measured every 4 hours over 24 hours. There were some missing measurement, but each participant had 4 ∼ 6 measurement, resulting in a total of 63 samples in restricted group and 62 samples in unrestricted group. Detailed description of this study has been previously published [35]. This RNA-seq dataset is publicly available in GEO (GES129843). After filtering the gene probes with mean cpm less than 1, 13,167 gene probes were kept. We further performed log2 transformation [i.e., log 2(*x* + 1), where *x* is the cpm of a gene in a sample] to improve the normality of the data.

### Circadian pattern detection

We first applied the LR rhythmicity method to this time-restricted feeding dataset. Under *p*_*c*_ *≤* 0.01, we identified 1,407 and 935 genes showing significant circadian rhythmicity for the restricted group and the unrestricted group, respectively. Figure S14 and S15 shows the 6 core circadian genes in the TRF group and the URF group, including *PER1, PER2, PER3, ARNTL, NR1D1*, and *DBP*, which are known to have persistent circadian rhythmicity. For these 6 circadian genes for the restricted group and the unrestricted group, our method (LR rhythmicity) yielded highly significant p-value (1.35× 10^−8^ ∼ 1.65 × 10^−21^), showing the strong detection power of circadian rhythmicity. We further performed pathway enrichment analysis. Using *p*_*pathway*_ *≤* 0.01 as cutoff, our likelihood methods detected 61 and 105 significant pathways for the TRF group and URF group respectively. The top pathways enriched in both groups included Circadian Rhythm Signaling pathway, Prolactin Signaling pathway, IGF-1 Signaling pathway, which are all known to related with circadian rhythmicity [36, 37].

### Differential circadian analysis

We further performed differential circadian analysis comparing TRF and URF groups using LR diff. In terms of differential fit, we started with candidate genes that showed circadian rhythmicity (*p*_*c*_ *≤* 0.01) in either restricted or unrestricted (n=1,864). Comparing the TRF group to the URF group (baseline group), LR diff identified 57 genes showing differential fit 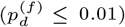. The most significant gene, *RUFY* 1, is shown in Figure S16(d), where there was a rhythmicity in the TRF group but not in the URF group.

In terms of differential amplitude, differential phase, and differential basal level, we started with candidate genes that showed circadian rhythmicity (*p*_*c*_ *≤* 0.01) in both TRF and URF (n=478). Comparing TRF to URF, 11 genes showing differential amplitude 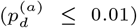, 25 genes showing differential phase 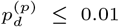, 8 genes showing differential basal level 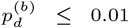. Figure S16(a-c) showed the most significant genes for differential amplitude, phase, and basal level comparing TRF and URF groups, respectively.

## Discussion

In summary, we developed a series of likelihood-based methods for detecting (i) circadian rhythmicity and (ii) differential circadian patterns. In terms of circadian rhythmicity detection, our method (LR rhythmicity) could better control the type I error rate to its nominal level (i.e., produce an accurate p-values) than the other competing methods. In terms of differential circadian patterns, our likelihood-based method is the first parametric method to characterize 4 sub-categories of differential circadian patterns, including differential amplitude, differential phase, differential basal level, and differential fit. Simulation shows that our method (LR diff) successfully controlled the type I error rate to the 5% nominal level for all 4 types of differential circadian patterns under the Gaussian assumption. In addition, LR diff was more powerful than the competing methods in terms of differential fit. We also applied our methods in transcriptomic data applications including a brain aging data, and a time restricted feeding data. Superior performance has been observed in these applications.

Our methods have the following strengths. (i) The type I error rate of both LR rhythmicity and LR diff were well controlled, indicating the p-values from these methods are accurate. While in the literature, it remained a concern about the type I error rate control for existing methods in terms of detecting circadian rhythmicity. (ii) Some methods require integer input circadian time, and even intervals between adjacent circadian time, while our methods have no such restrictions. Circadian time from modern epidemiology studies usually unevenly distributed between 0 hours and 24 hours. Thus, our method can be more applicable in biomedical applications. (iii) For examining differential fit, our method is statistically more powerful compared to other existing methods.

Our methods could potentially suffer from the following limitation. Our proposed methods are based on likelihood, which assume the residuals are normally distributed. And as shown in our simulation, the violation of such Gaussian assumptions may result in inflated type I error rate. To address this issue, we would recommend users to check the normality assumptions of the residuals. If the residuals (i.e., *y*_*i*_ − *ŷ*_*i*_) violated the Gaussian assumption, we would recommend to take transformation (i.e., log transformation) of the raw data to improve normality. We have adopted this approach for the dietary restriction real data example.

We plan to do the following future works. (i) In epidemiology studies, many other biological factors (e.g., age, gender, etc) could have a confounding impact on the circadian rhythmicity. Thus we will enable adjusting for covariates under our likelihood-based framework. (ii) To the best of our knowledge, no circadian rhythmicity detection method could handle repeated measurement from the same individuals. For example, the time restricted feeding data example employed a cross-over design, and the 11 participants with each participant repeatedly measured 4-6 times. By extending our methods to model this within subject correlation, we would expect higher power to detect circadian rhythmicity and differential circadian patterns.

An R package for our method is publicly available on GitHub https://github.com/diffCircadian/diffCircadian.

## Supporting information

diffCircadian_supplemental

## Acknowledgments

We thank the anonymous reviewers for their valuable suggestions.

