## Supplementary material for "Likelihood-based Tests for Detecting Circadian Rhythmicity and Differential Circadian Patterns in Transcriptomic Applications": diffCircadian_supplemental

*Corresponding author

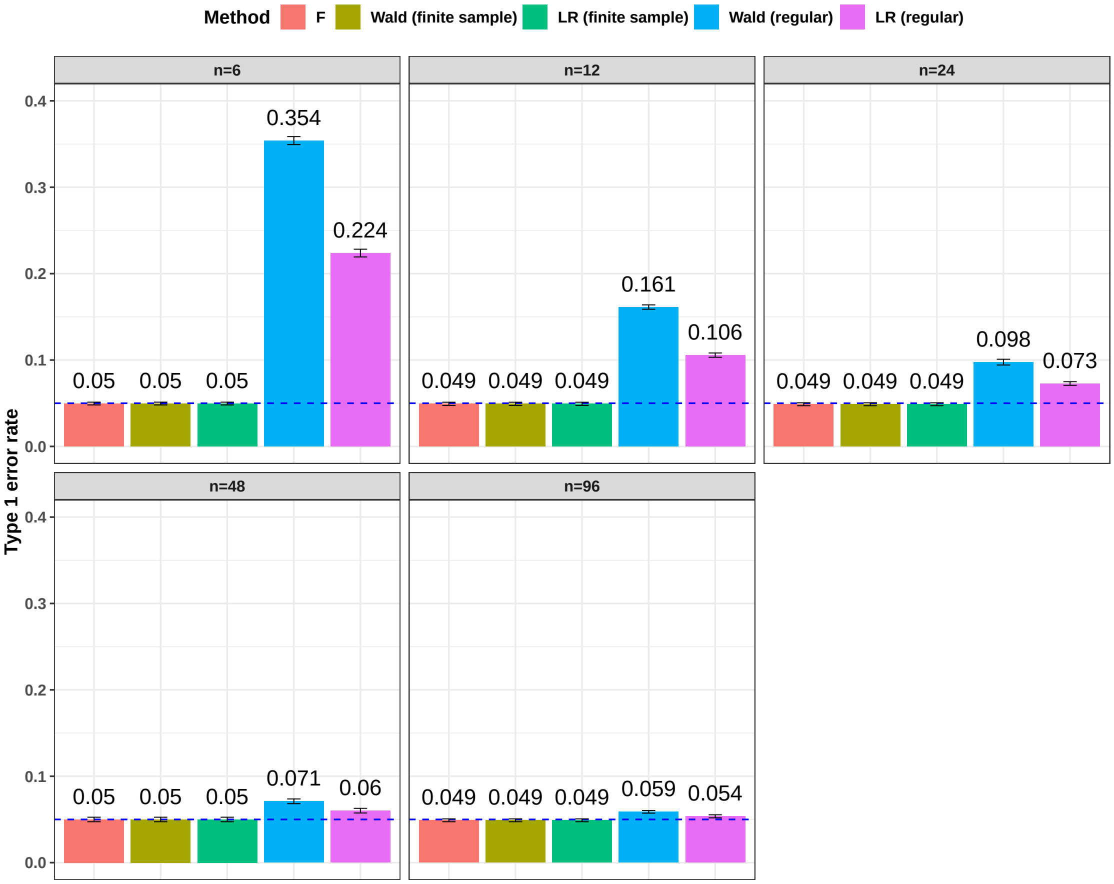


Figure S1: Type I error rate at nominal $\alpha$ level 5% for F test and the four likelihood-based methods in detecting circadian rhythmicity. The sample sizes were varied at n=6, 12, 24, 48, and 96. The blue dashed line is the 5% nominal level. A higher than 5% blue dashed line bar indicates an inflated type I error rate; a lower than 5% blue dashed line bar indicates a conservative type I error rate; and a bar at the blue dashed line indicates an accurate type I error rate (i.e., p-value = 0.05). The standard deviation of the mean type I error rate was also marked on the bar plot.

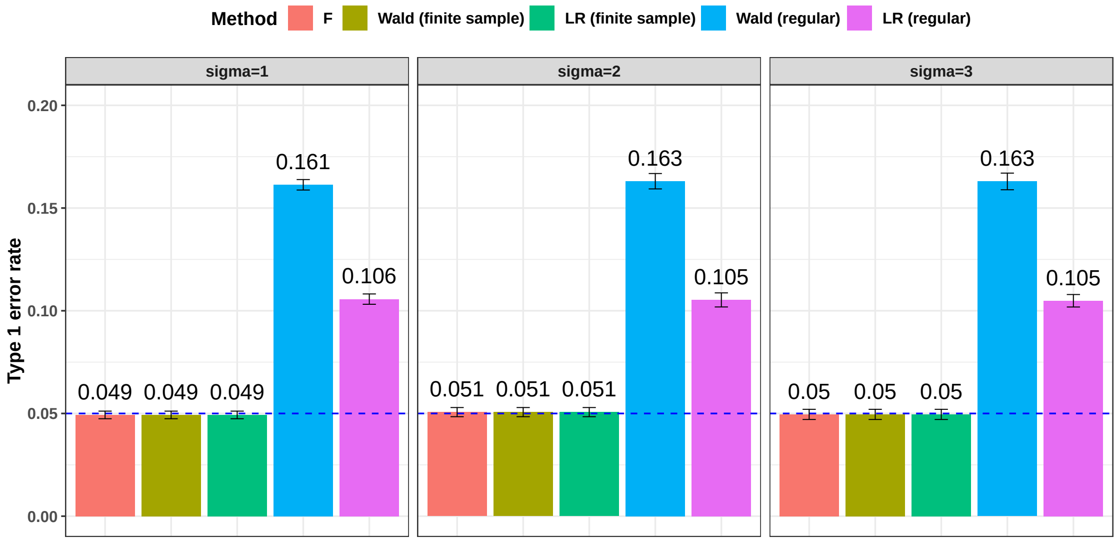


Figure S2: Type I error rate at nominal $\alpha$ level 5% for F test and the four likelihood-based methods in detecting circadian rhythmicity. The noise level were varied at $\sigma = 1, 2, 3$. The blue dashed line is the 5% nominal level. A higher than 5% blue dashed line bar indicates an inflated type I error rate; a lower than 5% blue dashed line bar indicates a conservative type I error rate; and a bar at the blue dashed line indicates an accurate type I error rate (i.e., p-value = 0.05). The standard deviation of the mean type I error rate was also marked on the bar plot.

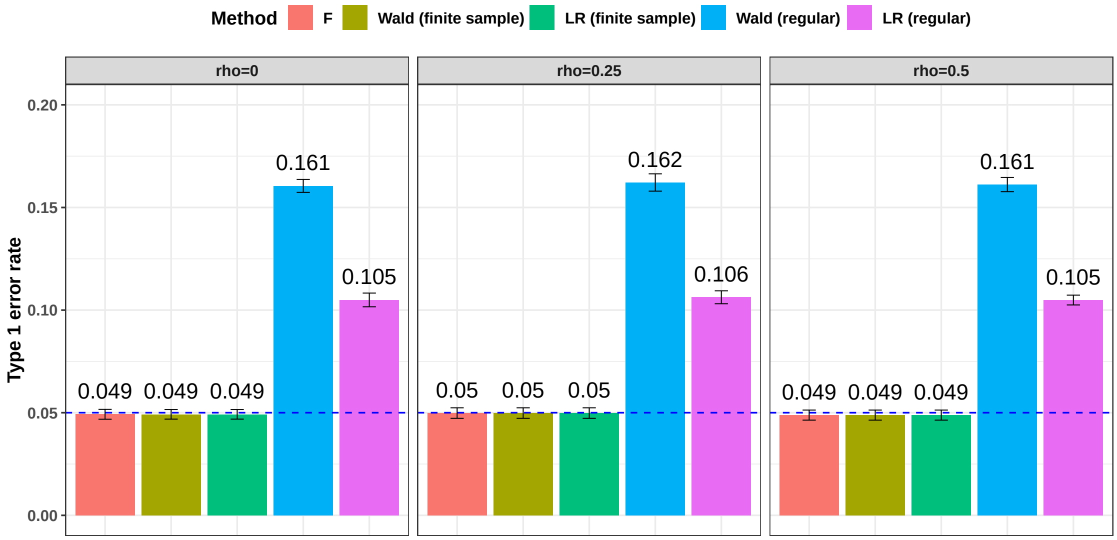


Figure S3: Type I error rate at nominal $\alpha$ level 5% for F test and the four likelihood-based methods in detecting circadian rhythmicity. The correlation strength between genes were varied at $\rho = 0, 0.25, 0.5$. The blue dashed line is the 5% nominal level. A higher than 5% blue dashed line bar indicates an inflated type I error rate; a lower than 5% blue dashed line bar indicates a conservative type I error rate; and a bar at the blue dashed line indicates an accurate type I error rate (i.e., p-value = 0.05). The standard deviation of the mean type I error rate was also marked on the bar plot.

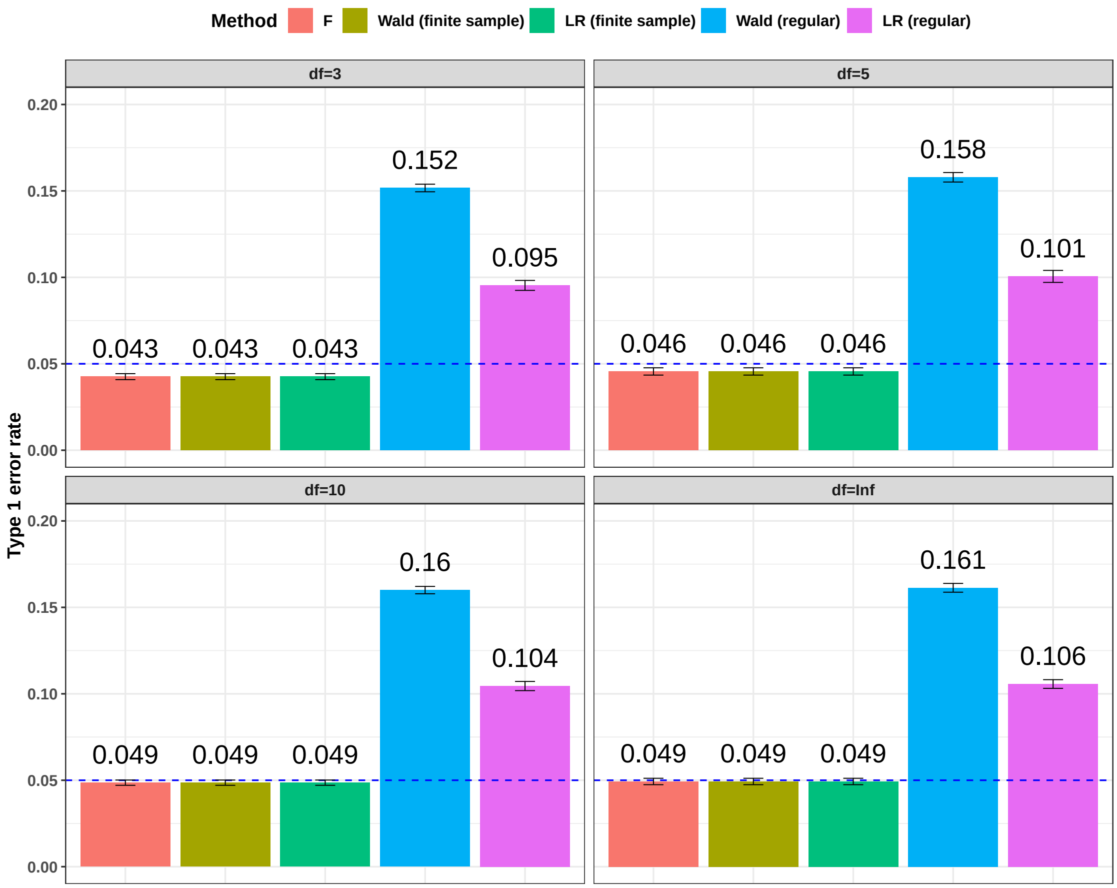


Figure S4: Type I error rate at nominal $\alpha$ level 5% for F test and the four likelihood-based methods in detecting circadian rhythmicity. The violation of the normality assumption $df$ were varied at $df = 3, 5, 10, \infty$, where $df$ is the degree of freedom of a t-distribution. When $df = \infty$, $t(\infty)$ is equivalent to a standard normal distribution (i.e., $N(0, 1)$). The blue dashed line is the 5% nominal level. A higher than 5% blue dashed line bar indicates an inflated type I error rate; a lower than 5% blue dashed line bar indicates a conservative type I error rate; and a bar at the blue dashed line indicates an accurate type I error rate (i.e., p-value = 0.05). The standard deviation of the mean type I error rate was also marked on the bar plot.

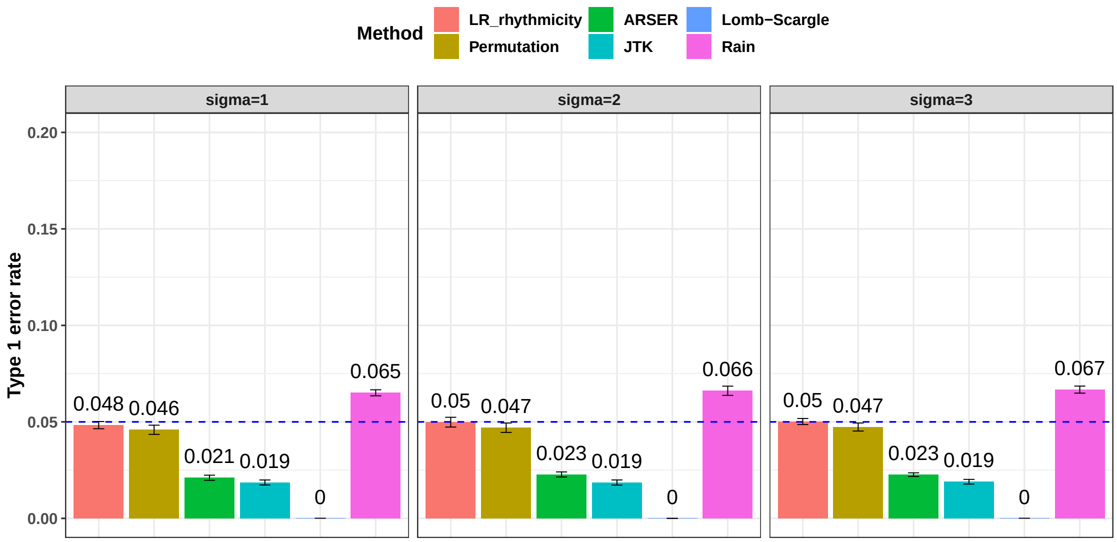


Figure S5: Type I error rate at nominal $\alpha$ level 5% for 6 different methods in detecting circadian rhythmicity. The noise level were varied at $\sigma = 1, 2, 3$. The blue dashed line is the 5% nominal level. A higher than 5% blue dashed line bar indicates an inflated type I error rate; a lower than 5% blue dashed line bar indicates a conservative type I error rate; and a bar at the blue dashed line indicates an accurate type I error rate (i.e., p-value = 0.05). The standard deviation of the mean type I error rate was also marked on the bar plot.

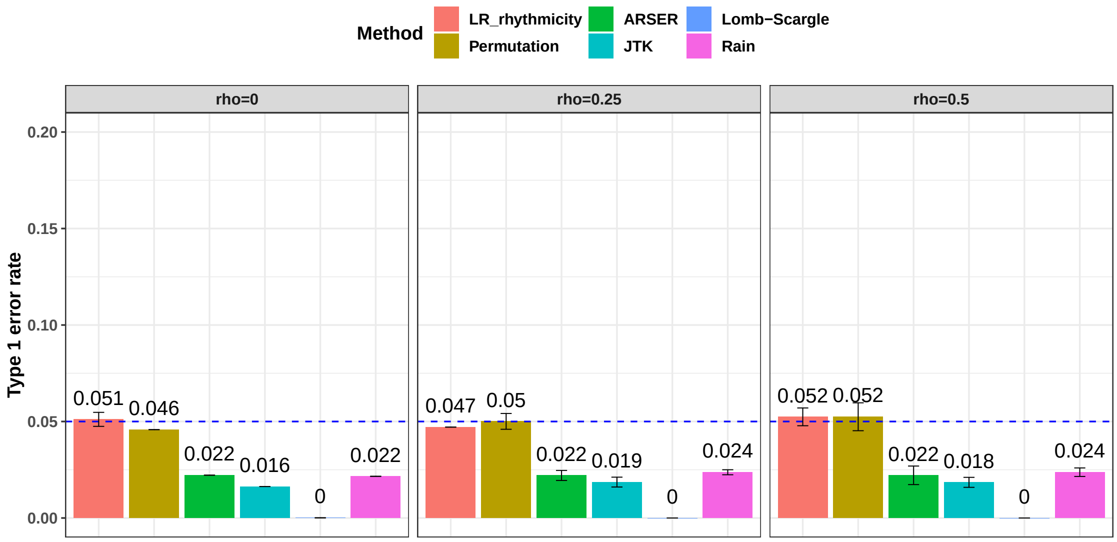


Figure S6: Type I error rate at nominal $\alpha$ level 5% for 6 different methods in detecting circadian rhythmicity. A higher than 5% blue dashed line bar indicates an inflated type I error rate; a lower than 5% blue dashed line bar indicates a conservative type I error rate; and a bar at the blue dashed line indicates an accurate type I error rate (i.e., p-value = 0.05). The correlation strength between genes were varied at $\rho = 0, 0.25, 0.5$. The standard deviation of the mean type I error rate was also marked on the bar plot.

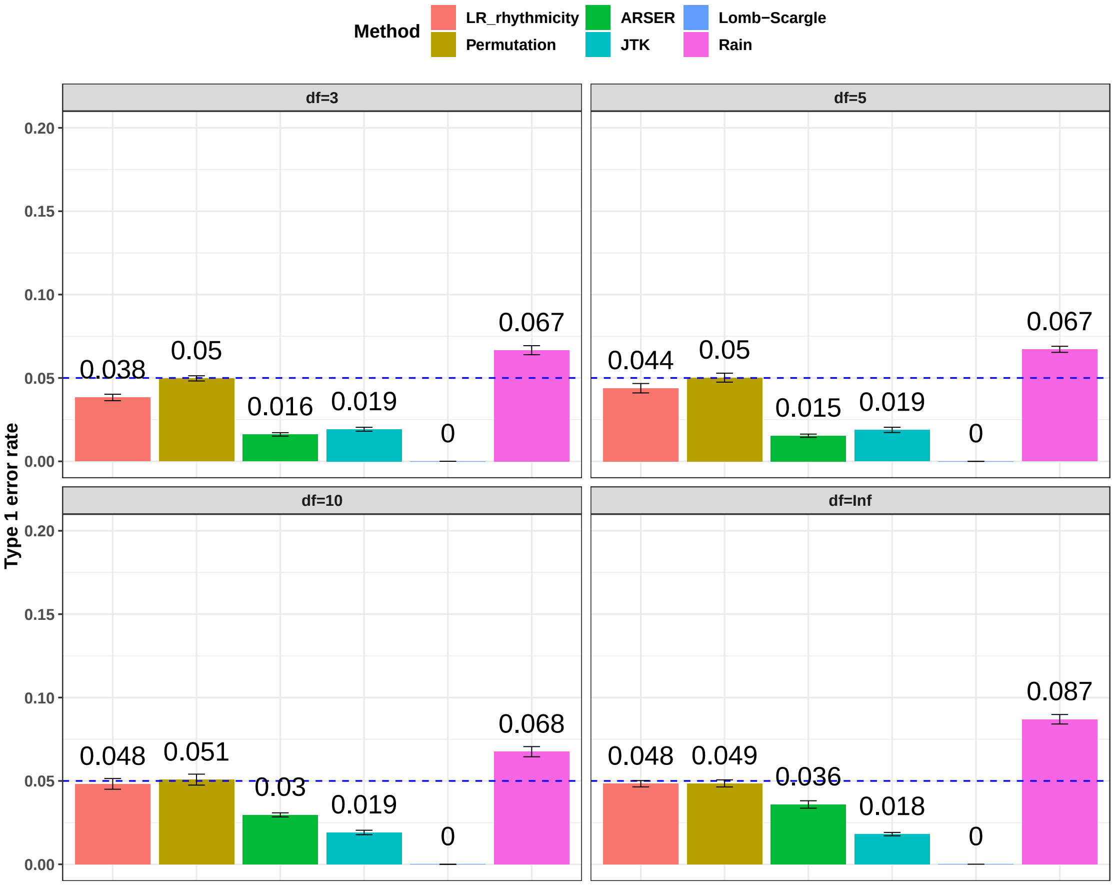


Figure S7: Type I error rate at nominal $\alpha$ level 5% for 6 different methods in detecting circadian rhythmicity. The violation of the normality assumption $df$ were varied at $df = 3, 5, 10, \infty$, where $df$ is the degree of freedom of a t-distribution. When $df = \infty$, $t(\infty)$ is equivalent to a standard normal distribution (i.e., $N(0, 1)$). The blue dashed line is the 5% nominal level. A higher than 5% blue dashed line bar indicates an inflated type I error rate; a lower than 5% blue dashed line bar indicates a conservative type I error rate; and a bar at the blue dashed line indicates an accurate type I error rate (i.e., p-value = 0.05). The standard deviation of the mean type I error rate was also marked on the bar plot.

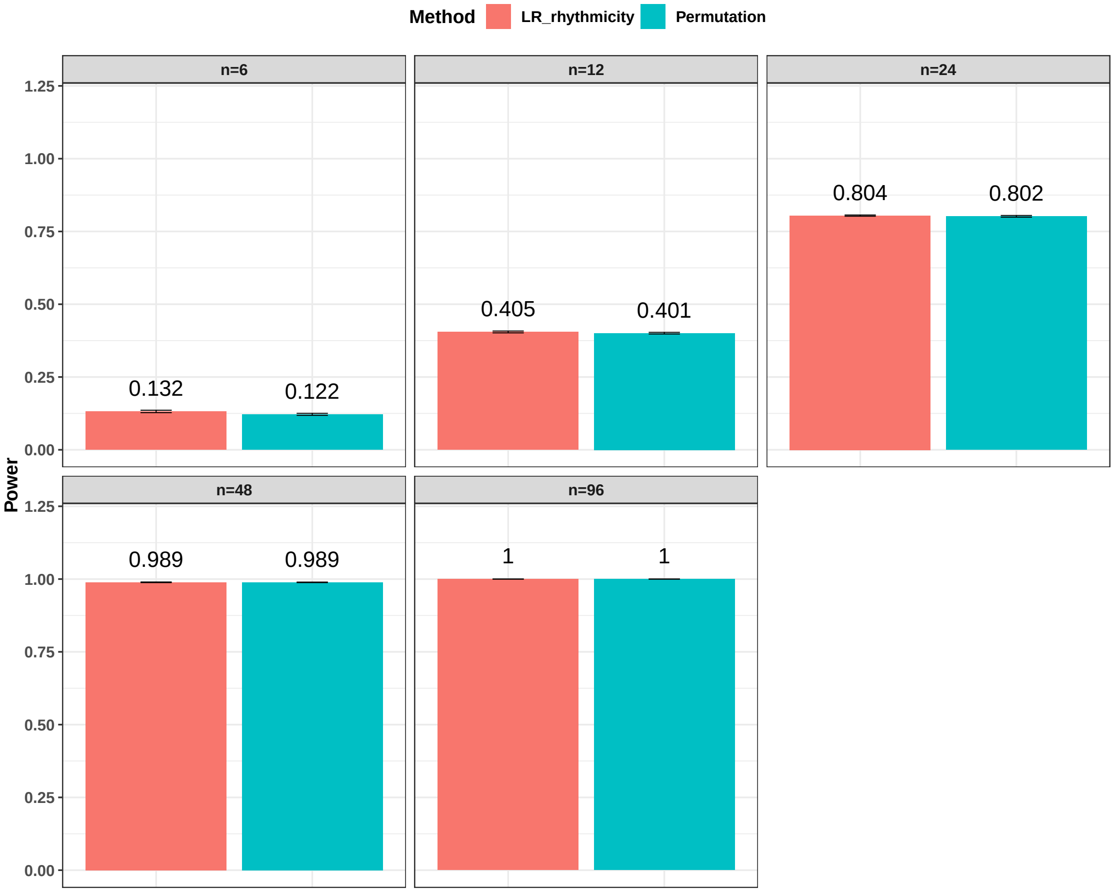


Figure S8: Power evaluation for LR_rhythmicity and the permutation test. The sample sizes were varied at n=6, 12, 24, 48, and 96. The standard deviation of the mean type I error rate was also marked on the bar plot.

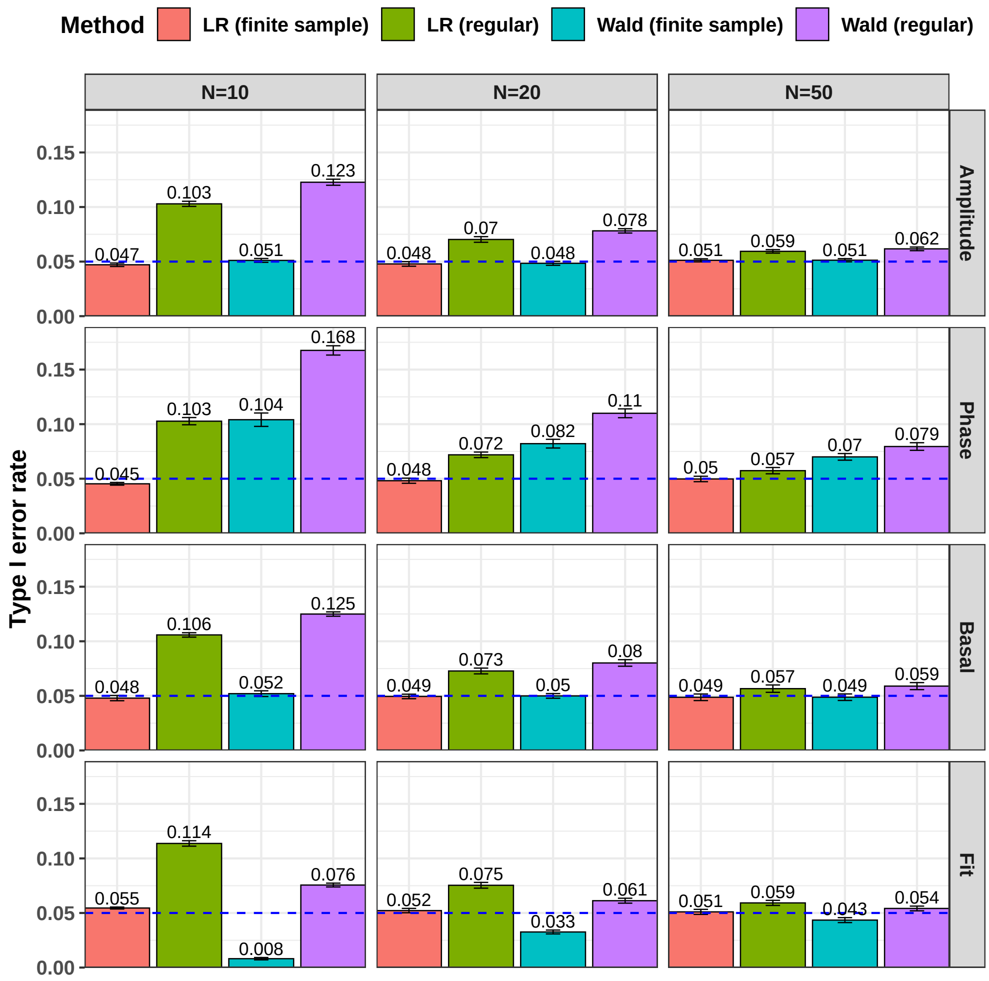


Figure S9: Type I error rate at nominal $\alpha$ level 5% for the 4 likelihood-based methods in detecting differential circadian patterns. The differential circadian patterns include differential amplitude (Amplitude), differential phase (Phase), differential basal level (Basal), and differential fit (Fit). The sample sizes were varied at N=10, 20, and 50. The blue dashed line is the 5% nominal level. A higher than 5% blue dashed line bar indicates an inflated type I error rate; a lower than 5% blue dashed line bar indicates a conservative type I error rate; and a bar at the blue dashed line indicates an accurate type I error rate (i.e., p-value = 0.05). The standard deviation of the mean type I error rate was also marked on the bar plot.

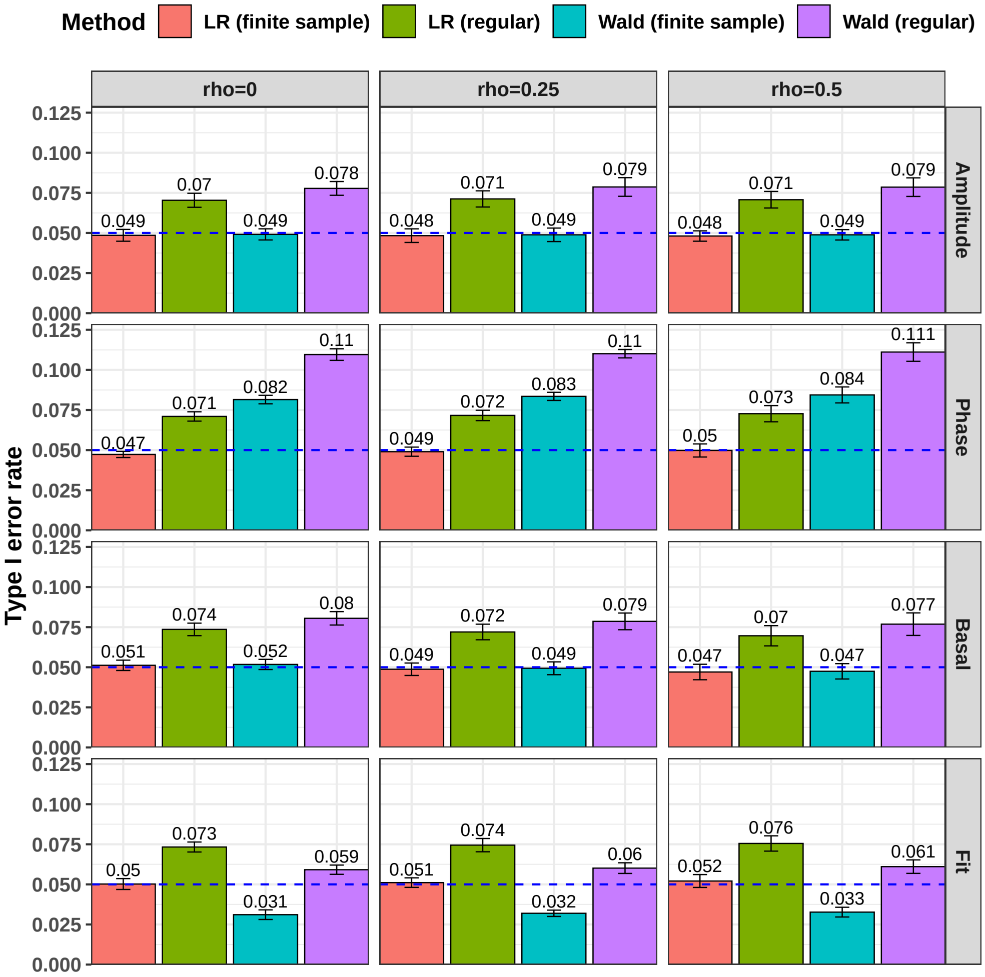


Figure S10: Type I error rate at nominal $\alpha$ level 5% for 6 different methods in detecting differential circadian patterns. The differential circadian patterns include differential amplitude (Amplitude), differential phase (Phase), differential basal level (Basal), and differential fit (Fit). The correlation strength between genes were varied at $\rho = 0, 0.25, 0.5$. The blue dashed line is the 5% nominal level. A higher than 5% blue dashed line bar indicates an inflated type I error rate; a lower than 5% blue dashed line bar indicates a conservative type I error rate; and a bar at the blue dashed line indicates an accurate type I error rate (i.e., p-value = 0.05). The standard deviation of the mean type I error rate was also marked on the bar plot.

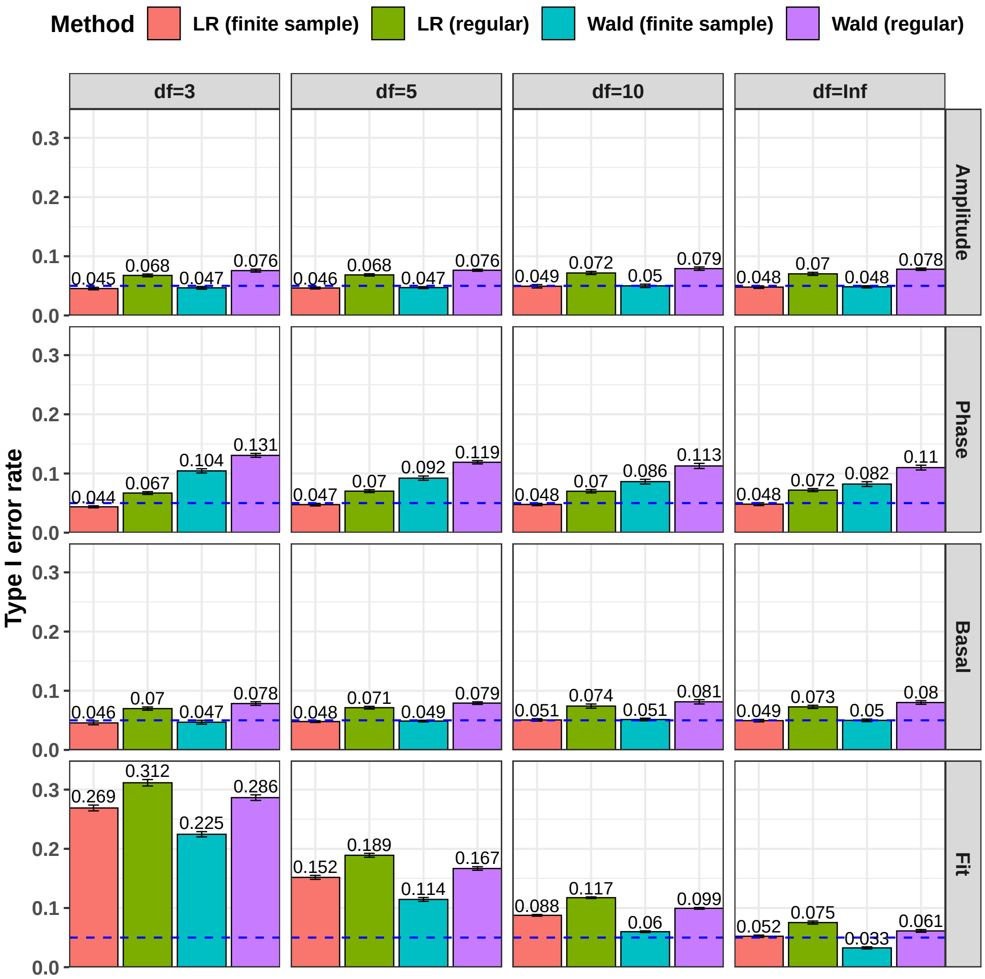


Figure S11: Type I error rate at nominal $\alpha$ level 5% for 6 different methods in detecting differential circadian patterns. The differential circadian patterns include differential amplitude (Amplitude), differential phase (Phase), differential basal level (Basal), and differential fit (Fit). The violation of the normality assumption $df$ were varied at $df = 3, 5, 10, \infty$, where $df$ is the degree of freedom of a t-distribution. When $df = \infty$, $t(\infty)$ is equivalent to a standard normal distribution (i.e., $N(0, 1)$). The blue dashed line is the 5% nominal level. A higher than 5% blue dashed line bar indicates an inflated type I error rate; a lower than 5% blue dashed line bar indicates a conservative type I error rate; and a bar at the blue dashed line indicates an accurate type I error rate (i.e., p-value = 0.05). The standard deviation of the mean type I error rate was also marked on the bar plot.

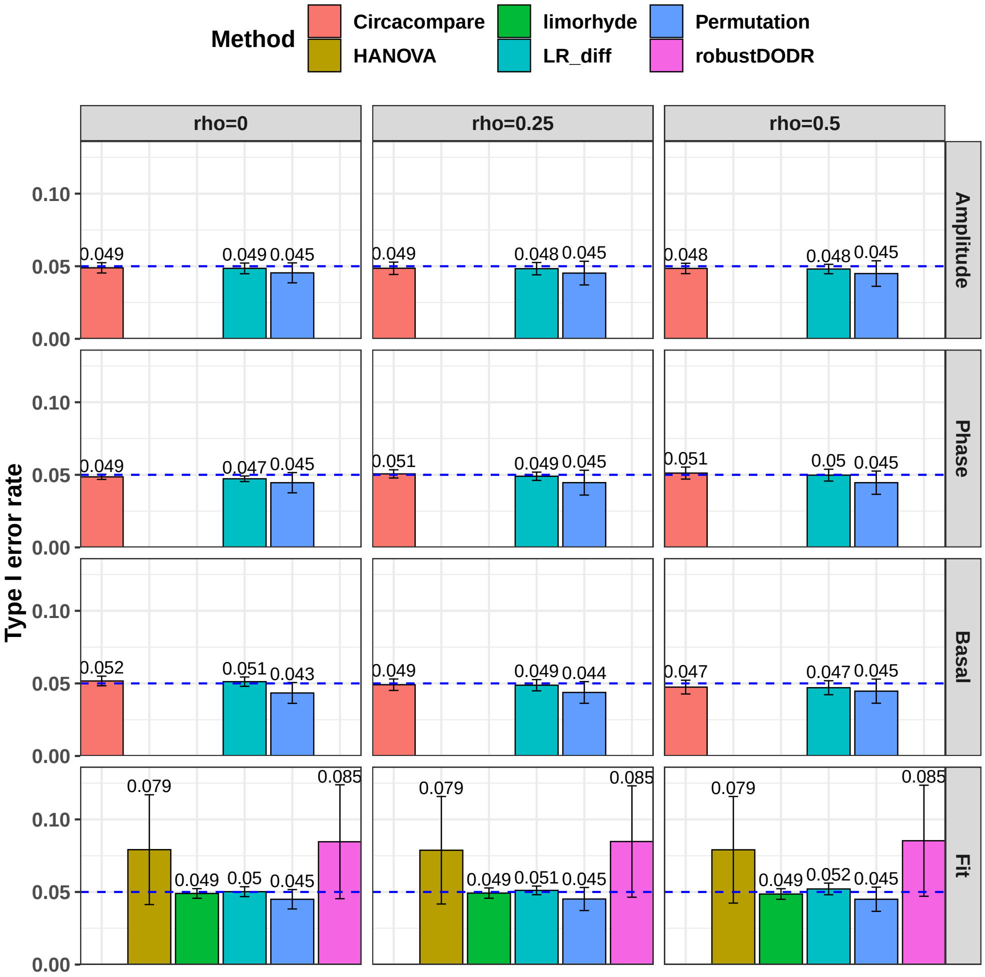


Figure S12: Type I error rate at nominal $\alpha$ level 5% for 6 different methods in detecting differential circadian patterns. The differential circadian patterns include differential amplitude (Amplitude), differential phase (Phase), differential basal level (Basal), and differential fit (Fit). The correlation strength between genes were varied at $\rho = 0, 0.25, 0.5$. The blue dashed line is the 5% nominal level. A higher than 5% blue dashed line bar indicates an inflated type I error rate; a lower than 5% blue dashed line bar indicates a conservative type I error rate; and a bar at the blue dashed line indicates an accurate type I error rate (i.e., p-value = 0.05). The standard deviation of the mean type I error rate was also marked on the bar plot.

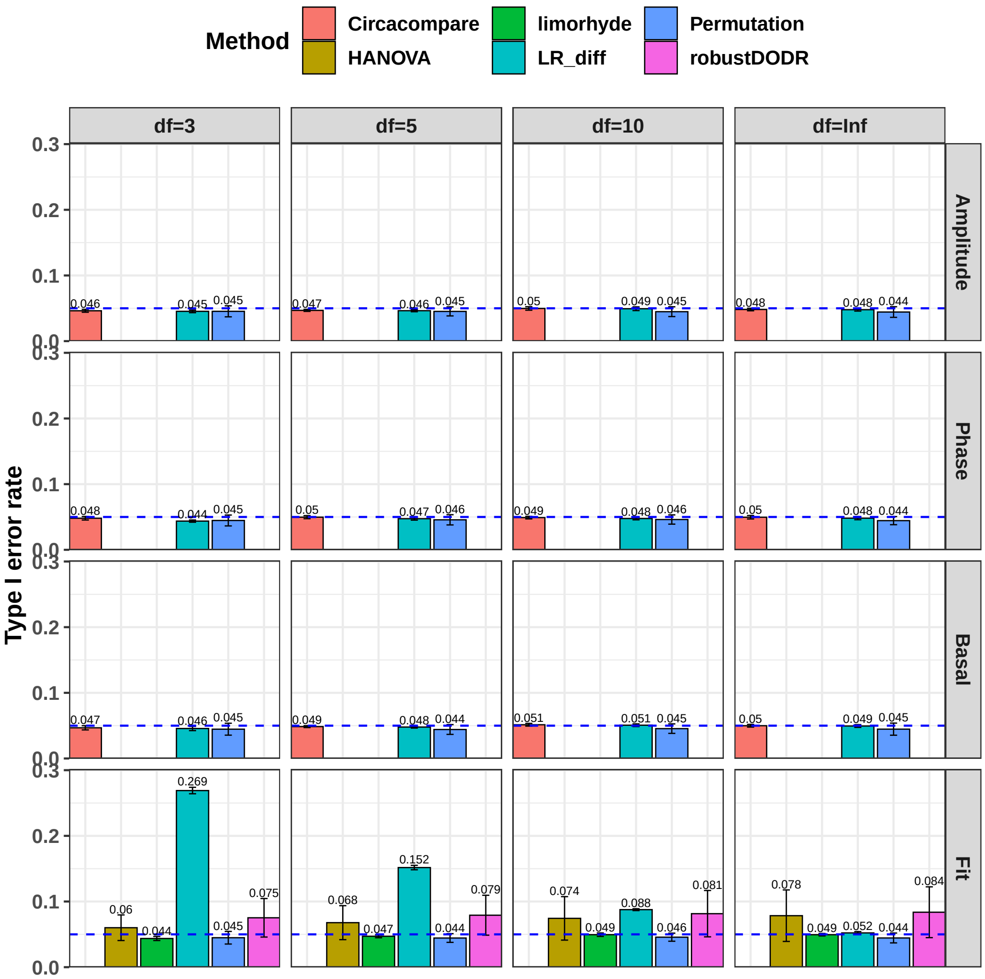


Figure S13: Type I error rate at nominal $\alpha$ level 5% for 6 different methods in detecting differential circadian patterns. The differential circadian patterns include differential amplitude (Amplitude), differential phase (Phase), differential basal level (Basal), and differential fit (Fit). The violation of the normality assumption $df$ were varied at $df = 3, 5, 10, \infty$, where $df$ is the degree of freedom of a t-distribution. When $df = \infty$, $t(\infty)$ is equivalent to a standard normal distribution (i.e., $N(0, 1)$). The blue dashed line is the 5% nominal level. A higher than 5% blue dashed line bar indicates an inflated type I error rate; a lower than 5% blue dashed line bar indicates a conservative type I error rate; and a bar at the blue dashed line indicates an accurate type I error rate (i.e., p-value = 0.05). The standard deviation of the mean type I error rate was also marked on the bar plot.

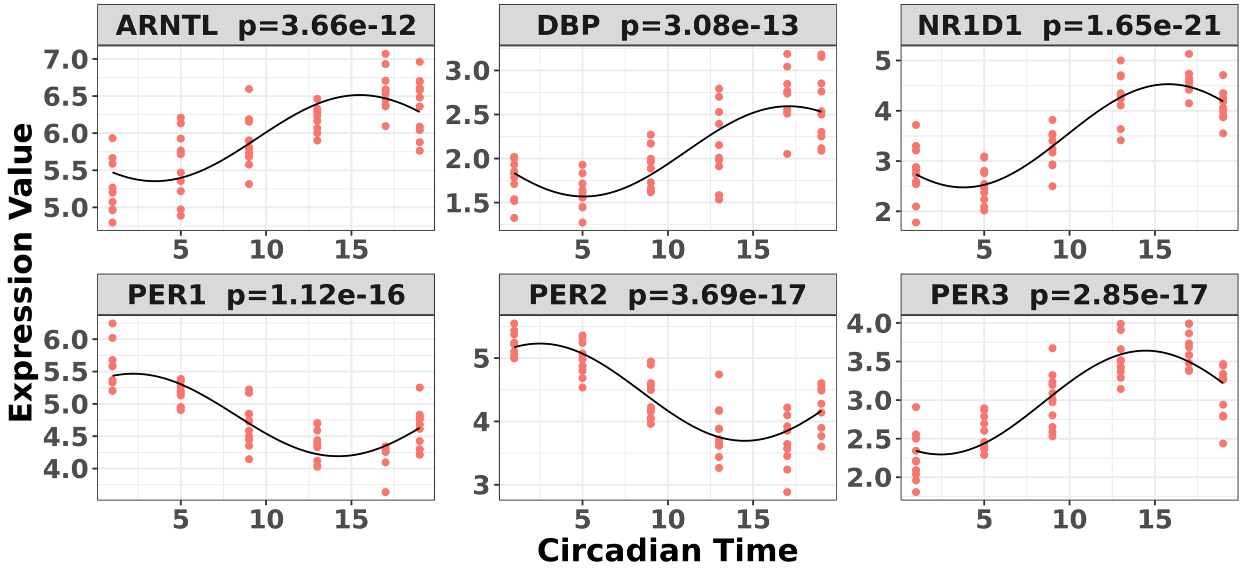


Figure S14: Circadian rhythmicity detected by LR_rhythmicity for 6 core circadian genes in the restricted group of the time-restricted feeding data, including PER1, PER2, PER3, ARNTL, NR1D1, and DBP.

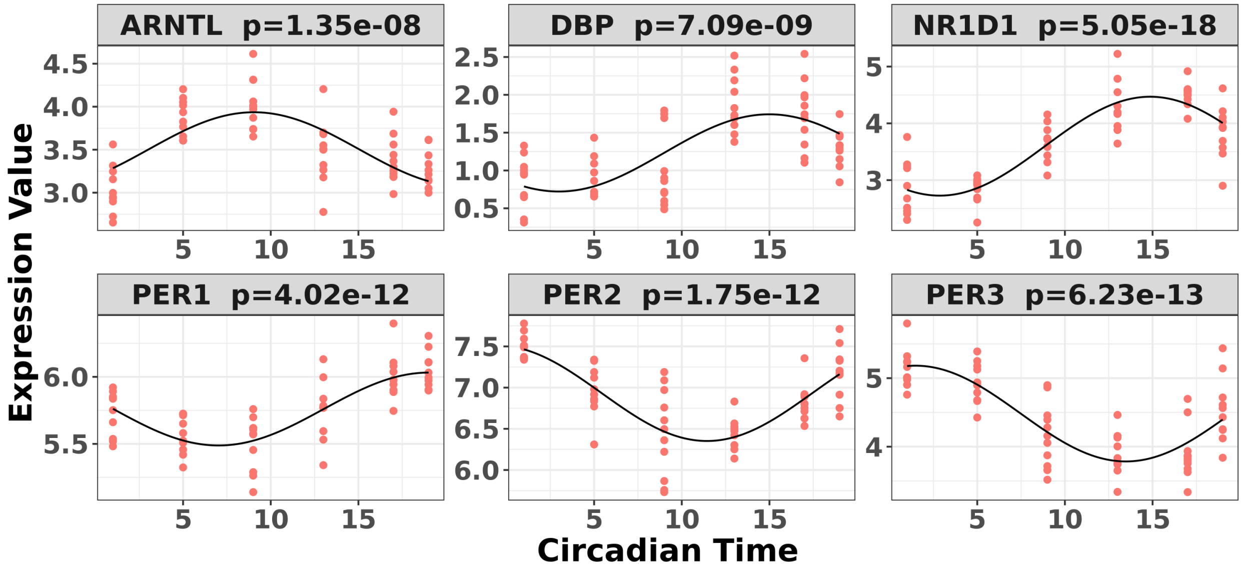


Figure S15: Circadian rhythmicity detected by LR_rhythmicity for 6 core circadian genes in the unrestricted group of the time-restricted feeding data, including PER1, PER2, PER3, ARNTL, NR1D1, and DBP.

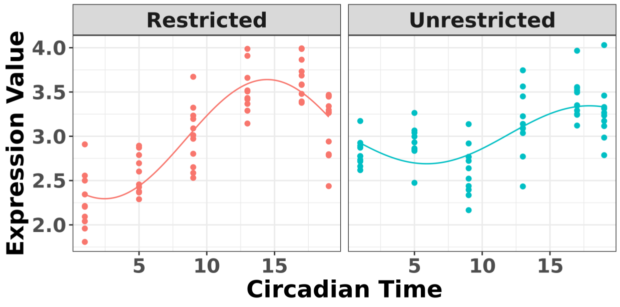


(a) Differential amplitude for KLHL25 ($p = 8.80 \times 10^{-6}$)

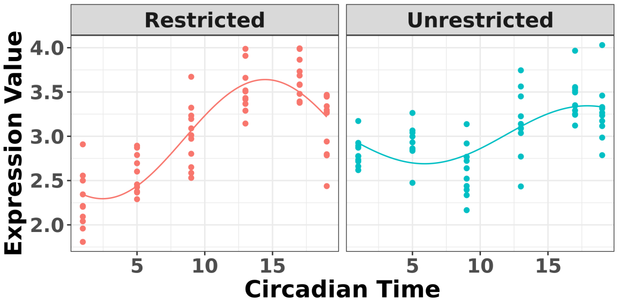


(b) Differential basal for KLHL25 ($p = 4.47 \times 10^{-5}$)

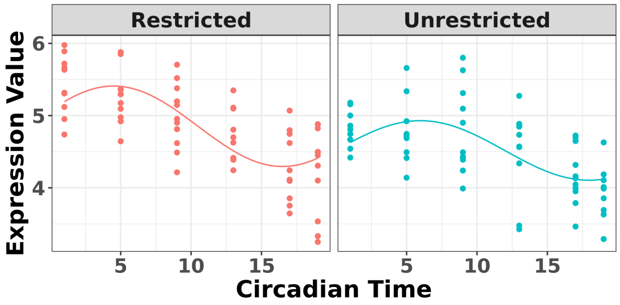


(c) Differential phase for TSPAN8 ($p = 1.37 \times 10^{-4}$)

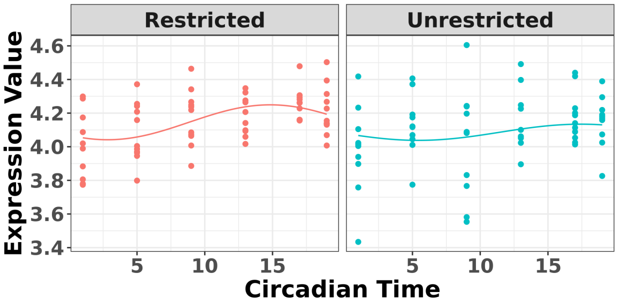


(d) Differential fit for RUFY1 ($p = 7.70 \times 10^{-5}$)

Figure S16: The most significant genes showing four types of differential circadian patterns from the time-restricted feeding data.
